# Acetylcholine engages distinct amygdala microcircuits to gate internal theta rhythm

**DOI:** 10.1101/2023.02.26.530135

**Authors:** Joshua X. Bratsch-Prince, James W. Warren, Grace C. Jones, Alexander J. McDonald, David D. Mott

## Abstract

Acetylcholine (ACh) is released from basal forebrain cholinergic neurons in response to salient stimuli and engages brain states supporting attention and memory. These high ACh states are associated with theta oscillations, which synchronize neuronal ensembles. Theta oscillations in basolateral amygdala (BLA) underlie emotional memory, yet their mechanism remains unclear. Using brain slice electrophysiology in mice, we show large ACh stimuli evoke prolonged theta oscillations in BLA local field potential that depend upon activation of cholecystokinin (CCK) interneurons (INs). Somatostatin (SOM) INs inhibit CCK INs and are themselves inhibited by ACh, gating BLA theta. ACh-induced theta activity is more readily evoked in BLA over cortex or hippocampus, suggesting preferential activation of BLA during high ACh states. These data reveal a SOM-CCK IN circuit in BLA that gates internal theta oscillations and suggest a mechanism by which salient stimuli acting through ACh switch the BLA into a network state enabling emotional memory.

## Introduction

Alterations in behavioral state are associated with changes in brain activity. Subcortical neuromodulators are instrumental in regulating these state-dependent changes^1–5^. One such modulator is acetylcholine (ACh), which plays an important role in establishing network states that support attention, memory and emotional processes^6–10^. Cholinergic innervation of the cortex and limbic regions arises from the basal forebrain (BF)^11–14^. Cholinergic neurons in BF fire in response to emotionally salient stimuli and release ACh, promoting synaptic plasticity and providing favorable conditions for information encoding^15–18^. These high ACh brain states are associated with network oscillations in the theta frequency (3-12 Hz)^7,19–21^. Theta oscillations synchronize neuronal activity and allow binding of cell assemblies between brain regions. Interestingly, the basolateral amygdala (BLA) receives the most robust cholinergic innervation of any target of the cholinergic BF^22,23^, suggesting that ACh plays a central role in regulating BLA function. In the BLA of both humans and rodents theta oscillations are increased during emotional processing^24–28^ where they are thought to be a neural correlate of avoidance and freezing behavior^29,30^. BLA theta oscillations are synchronized with activity in both the ventral hippocampus (vHPC) and prelimbic cortex (PL) during emotional behaviors^25,28,31,32^, where the BLA transmits valence-related information to modulate avoidance and learned fear^33–35^. However, while ACh is critical in regulating these emotional behaviors, there is not a clear understanding of how it alters ensemble neuronal activity to promote theta oscillations.

Studies in hippocampus and cortex have found that ACh actions on GABAergic inhibitory interneurons (INs) are critical in regulating network oscillations^36–39^. GABAergic INs exhibit diverse anatomical and functional properties that endow them with distinct roles in generating local oscillations. In the BLA three largely non-overlapping IN populations can be divided based on those expressing parvalbumin (PV), cholecystokinin (CCK), and somatostatin (SOM)^40^. Anatomically, SOM INs target distal dendrites^41^, while PV and CCK INs perisomatically innervate local BLA excitatory pyramidal neurons (PYRs)^42,43^, forming independent inhibitory networks^44^. BLA INs are differently recruited during behavior to modulate BLA output^45,46^, but their precise roles in synchronizing BLA network activity are not well understood. Mechanistically, BLA INs could synchronize network activity through entrainment by rhythmic inputs from external regions or conversely, these rhythms could arise internally from local BLA circuit interactions. It is likely that both mechanisms occur and conditions favoring either are brain state specific. PV INs in BLA are strongly driven by glutamatergic input, providing a possible mechanism for external glutamatergic control of theta oscillations^47^. Indeed, PV INs have been implicated in BLA theta oscillations in some^48–50^, but not all conditions^51^. Alternately, ACh generates theta oscillation in the BLA *in vivo*, but also inhibits glutamate release from afferent cortical input^52^. This suggests favorable conditions for internally generated BLA oscillations. Whether local BLA circuit interactions can internally produce theta oscillations, however, is still not clear. Together. these observations highlight multiple possible circuit mechanisms for theta oscillations in the BLA. A comprehensive, cell-type-specific circuit understanding of ACh actions on theta oscillations in the BLA is needed to better understand the neural mechanisms underlying emotional behaviors.

In this study we systematically explored the role of BF derived ACh in generating theta oscillations in the BLA and the amygdalar circuits involved. We report that the effects of endogenous ACh on circuit dynamics are stimulus and brain area specific and identify a detailed mechanism by which ACh stimuli induce a network state driving BLA theta oscillations. Phasic cholinergic stimuli to the BLA act through muscarinic receptors (mAChRs) to reorganize local inhibitory circuits and produce internal theta oscillations with increased and synchronized PYR output. Both CCK and SOM INs play critical roles in this process. In particular, CCK INs are uniquely sensitive to ACh and are the drivers of this synchronized network behavior, revealing a previously undefined role for these cells in the BLA. These findings suggest a mechanism by which ACh release produced by emotionally salient stimuli can switch the BLA into a network state enabling emotional behavior and memory.

## Results

### Released acetylcholine bidirectionally modulates BLA theta oscillations

To explore how endogenously released ACh modulates BLA network activity, we used a double transgenic strategy to selectively express channelrhodopsin (ChR2) in choline acetyltransferase-expressing (ChAT) neurons (termed ChAT-ChR2 mice). These mice showed selective expression of ChR2 in cholinergic neurons in the basal forebrain and dense axonal expression of ChR2 in the BLA (Figure 1A).

**Figure 1.**
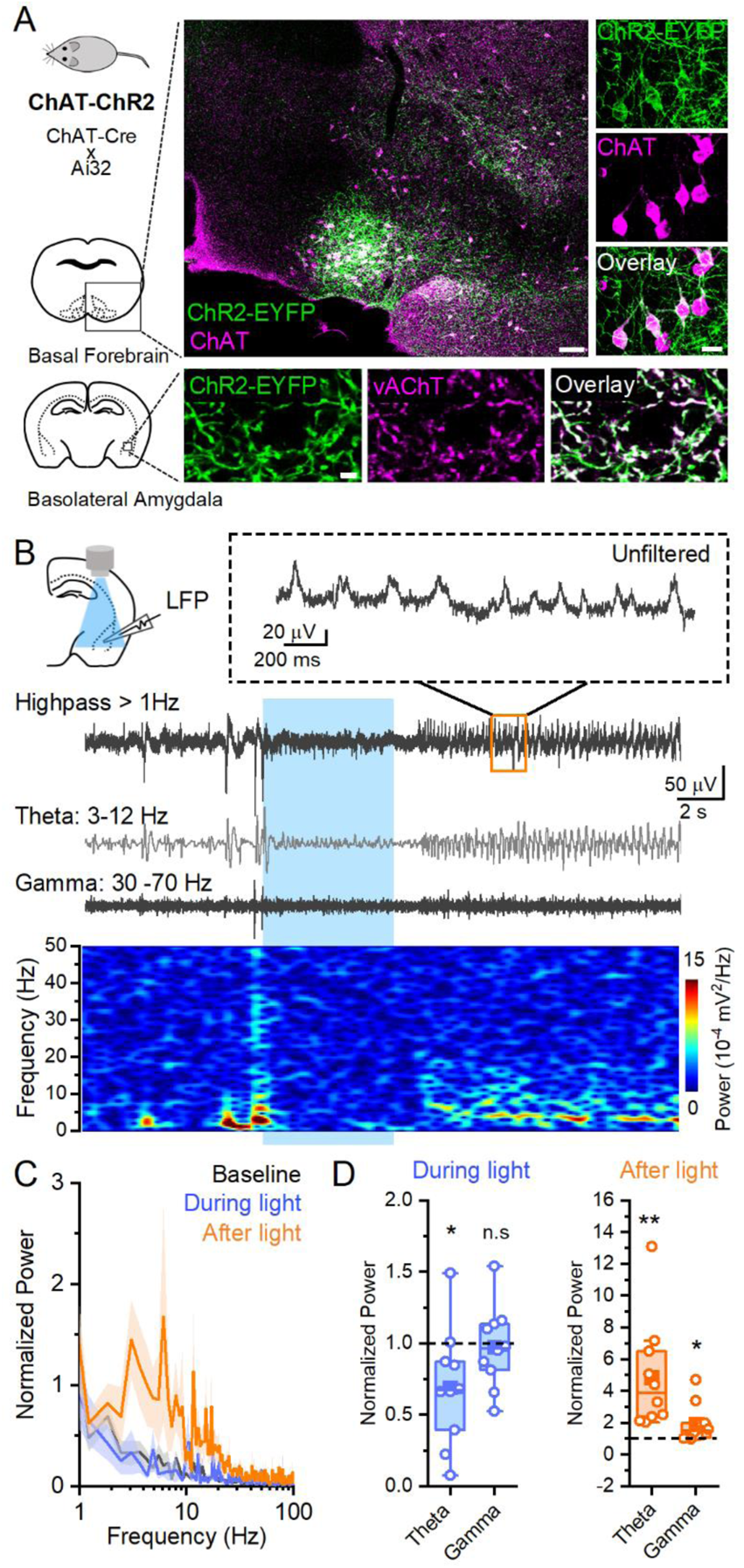
Optogenetically released acetylcholine bidirectionally modulates BLA LFP network activity. **A**. Confocal images of the basal forebrain and BLA in ChAT-ChR2 mice used for this study show ChAT expression in ChR2-expressing cells in the basal forebrain and vAChT expression in ChR2-expressing axons in the BLA. (99% of ChR2-EGFP+ also ChAT+, 79% of ChAT+ cells also ChR2-EGFP+; 877 cells, 2 animals). Scale bars 150, 25, and 5 μm. **B**. A representative waveform of BLA LFP recording showing a bidirectional effect of light stimulation (5 Hz for 5 s) on LFP activity in different frequency bands as separated with bandpass filters (short-time Fourier transform, below). **C**. Averaged normalized power spectrums from all slices before, during, and after light stimulation (10 slices (n), 6 animals (N)). **D**. During the light stimulus (left), the total power in theta frequency (3-12 Hz) is decreased (0.69 ± 0.12 of baseline; paired t-test, p = 0.039) while gamma frequency power (30-70 Hz) is not affected (0.97 ± 0.09 of baseline, paired t-test, p = 0.77). After light (right), both theta (4.85 ± 1.08 of baseline; paired t-test, p = 0.006) and gamma frequency power (1.92 ± 0.38 of baseline, paired t-test, p = 0.037) are increased.

Local field potential (LFP) recordings were performed in mouse coronal brain slices containing BLA. Cholinergic terminals were activated with 1-2 ms pulses of 490nm LED light delivered at 5 Hz for 5 seconds^53^. This stimulus pattern is consistent with basal forebrain cholinergic neuron firing *in vivo* during active waking and paradoxical sleep^54^, representing a physiological phasic ACh release event. Light stimulation of cholinergic terminals had a biphasic effect on BLA LFP activity. Low frequency activity in baseline was suppressed upon light stimulation that persisted during most of the 5-second light stimulus train (Figure 1B). Following light stimulation there was a robust increase in large amplitude rhythmic network activity in the theta range (3-12Hz) that often lasted for more than 10 seconds beyond the light stimulus. Power spectrum analysis in 2 second time bins before, during, and after light stimulation (Figure 1D) revealed that total power at theta and gamma frequencies were affected. During the light stimulus, total theta power was significantly suppressed while gamma power was not. After light, there was an almost five-fold increase in theta power and a smaller, but significant increase in gamma power. Comparing the magnitude of ACh increases in theta and gamma power showed the post-stimulation increase was significantly higher for the theta band (paired t-test, p = 0.025), highlighting a strong role of ACh in inducing theta oscillations in BLA slices, as is observed *in vivo*^55,56^.

### Acetylcholine increases inhibitory circuit activity in the BLA

To explore the circuit mechanisms underlying ACh’s induction of BLA theta oscillations, we performed intracellular whole cell recordings from BLA PYRs to monitor spontaneous synaptic activity. At baseline, recordings from PYRs showed periodic (< 0.5 Hz) large, compound events (Figure 2A). These events mirrored high amplitude, low frequency baseline activity of LFP recordings (Figure 1B) and were blocked by application of either glutamate receptor antagonists CNQX (20 μM) and APV (50 μM) or the GABA_A_ receptor antagonist bicuculline (20 μM), indicating they were dependent on both glutamate and GABA transmission, consistent with previous reports of low frequency network activity in BLA slices^57–59^. Light stimulation of cholinergic fibers induced a marked change in synaptic activity (Figure 2B). After an immediate (< 5 ms onset) large amplitude event induced by the first light pulse, there was a delayed induction of large, rhythmic events that occurred at theta frequency (Figure 2B, power spectrum analysis: peak power frequency = 6.93 ± 0.38 Hz: n = 51, N = 18) that persisted beyond the light stimulation (average duration = 10.24 ± 0.5 seconds). There was no difference in these events from slices of male or female mice, and both sexes were combined for the remainder of the study (Supplemental Table 1). Unlike large events in baseline, these theta frequency events were not affected by glutamate antagonists or by a GABA_B_ receptor antagonist but were fully blocked by bicuculline (Figure 2C), indicating these events were entirely GABAergic in nature and did not require glutamate transmission. These properties differed from low frequency BLA network activity in baseline states and highlight a switch in synchronized network dynamics by ACh. Additionally, while the broad nicotinic receptor (nAChR) antagonist mecamylamine (mec, 10 μM) had no effect on these events, they were fully blocked by the muscarinic receptor (mAChR) antagonist atropine (atro, 5 μM, Figure 2D), indicating they were driven by mAChRs, but not nAChRs. Further, application of an M1 (telenzepine (tzp), 0.1 μM) or an M2 (AFDX, 1 μM) antagonist had no effect on these events, but application of an M3 receptor antagonist (4DAMP, 1 μM) blocked them completely (Figure 2D). These results suggest ACh acts through an M3 receptor-mediated mechanism to drive inhibitory network activity in the BLA.

**Figure 2.**
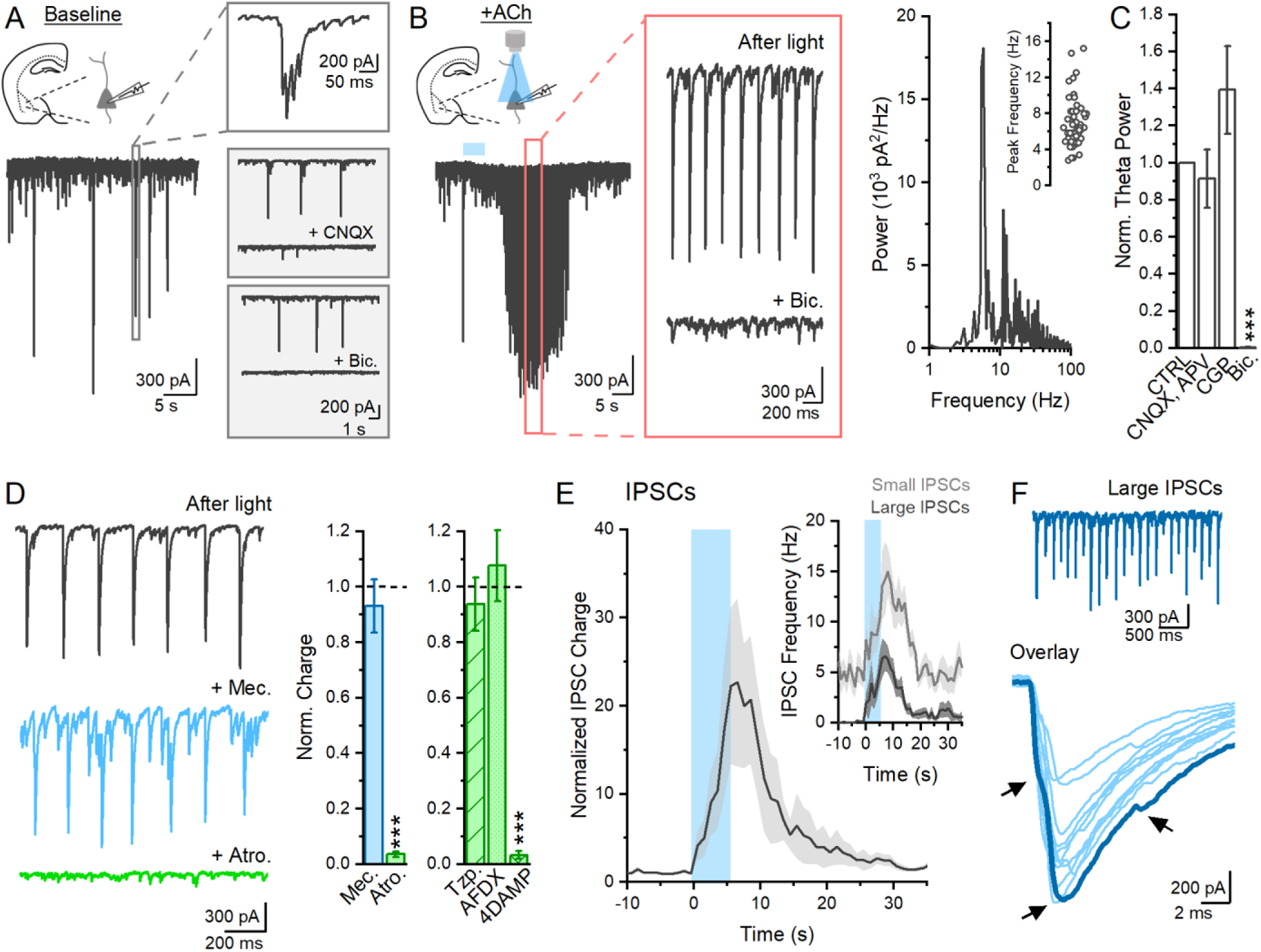
Acetylcholine induces large changes in BLA inhibitory activity. **A**. Baseline recordings of spontaneous network activity in BLA PYRs show low frequency, large amplitude events that are blocked by either CNQX (20 μM) or bicuculline (20 μM). **B**. Optogenetic stimulation (5 Hz, 5s) produces large theta frequency rhythmic events (power spectrum for sample waveform) that persist beyond light stimulation and are blocked by bicuculline (inset). **C**. Comparing total theta power of ACh-induced events show they are not blocked by glutamate antagonists (CNQX, 20 μM + APV, 50 μM, n = 12, N = 5) or a GABA_B_ antagonist (CGP, 2 μM, n = 6, N = 3) but are blocked by a GABA_A_ receptor antagonist (bicuculline, n = 7, N = 5), one way ANOVA, p=2.28×10^−5^. **D**. Sample waveforms from a representative cell showing rhythmic events after light in control (black), +Mec. (10 uM, blue), and +Atro. (5 uM, green). Normalized total charge of these events show they are not affected by Mec. but are blocked by Atro, n = 8, N = 6, one-way repeated measures ANOVA, p=1.66×10^−10^. They are also blocked by a selective M3 receptor antagonist 4DAMP (n = 8, N = 3), but not an M1 (Tzp, n = 5, N = 3) or M2 (AFDX, n = 8, N = 3) antagonist (green graph, right), one-way ANOVA, p=5.8×10^−10^. **E**. Isolation of IPSCs (CNQX and APV) following light stimulation shows a slow, robust increased in IPSC total charge and (inset) IPSC frequency of both small (1-3x average) and large (>5x average) amplitude IPSCs (n = 11, N = 6). **F**. Waveform from example cell showing large IPSCs (top). Superimposing each large IPSC shows they consistently contain multiple, summated IPCSs (arrows, example IPSC dark blue trace, other IPSCs light blue traces).

To understand the direct effects of ACh on BLA inhibitory activity, inhibitory post-synaptic currents (IPSCs) were isolated with application of glutamate antagonists (CNQX, 20 μM and APV, 50 μM). Measuring total IPSC charge over time (1 second bins) showed a drastic increase in IPSC charge in response to light stimulation that reached a maximal level after the end of light stimulation (Figure 2E). The increase in IPSC charge was not only due to the presence of larger amplitude theta IPSCs induced by light stimulation, but also an increase in overall IPSC frequency. Dividing IPSCs into small (1-3x average baseline IPSC amplitude) and large (>5x average baseline IPSC amplitude) events showed that light stimulation increased both types of IPSCs (Figure 2E), indicating an increase in overall inhibition in the BLA. Large IPSCs were significantly more enhanced by light stimulation and peaked in the theta frequency range (fold increase in IPSC frequency by light: large = 28.26 ± 4.36; small = 3.37 ± 0.43; paired t-test, p = 3.289 x 10^−4^). Isolation of the large IPSCs revealed that they contained tightly summated IPSCs that occurred within ms of each other (Figure 2F), suggesting highly synchronized activity from multiple presynaptic INs.

### Preferential activation of BLA over hippocampal or cortical inhibitory networks by acetylcholine

Due to the delayed timing of the rhythmic activity following stimulation, we were curious to explore how different amounts of cholinergic stimulation would impact circuit activity. Accordingly, increasing light stimulation (1, 5, 25, and 50 pulses at 5 Hz) was given either with or without the presence of low levels of the acetylcholinesterase inhibitor physostigmine (1 uM). Interestingly, in most cells, large rhythmic IPSCs were only induced with increasing cholinergic stimulation (Figure 3A) and rarely occurred with a single light pulse, suggesting they were dependent on the size of the cholinergic stimulus. In agreement, application of a low level of physostigmine greatly increased the probability of a given cholinergic stimulus to evoke large theta frequency IPSCs (Figure 3C). Comparing the frequency of evoked large IPSCs following light showed no difference across the increasing stimulations (Supplemental Figure 1), suggesting theta network activity in the BLA is evoked by large ACh stimuli independent of their nature. In cells that showed ACh-induced large IPSCs in ACSF control conditions, physostigmine did not affect the overall frequency of the large IPSCs but instead increased the time over which they occurred (Supplemental Figure 1), suggesting it did not fundamentally change the nature of the IPSCs and network event.

**Figure 3.**
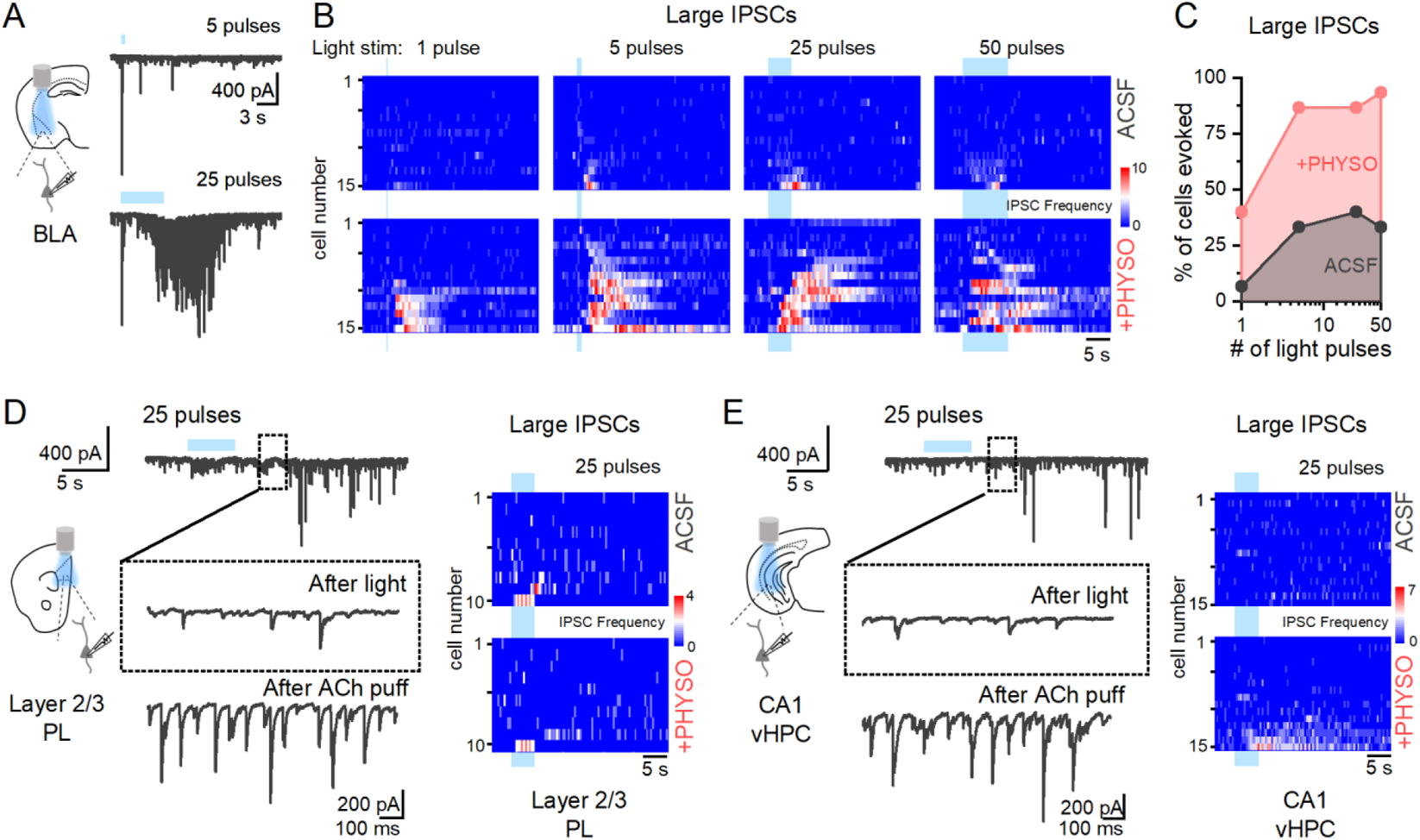
Optogenetically released acetylcholine more reliably activates BLA over PL or vHPC inhibitory networks. **A**. Sample recording from a BLA PYR where rhythmic theta activity was produced following 25 pulses of light stimulation (5 Hz), but not 5 pulses. **B.** Heatmaps showing large IPSC frequency (1s time bins) in all BLA cells that received 1, 5, 25, and 50 pulses of light stimulation (5 Hz) in ACSF (top) and in the presence of physostigmine (1uM). **C.** Plot showing the percentage of BLA cells that show sustained (> 1s) theta frequency large IPSCs in ACSF and physostigmine (n = 16, N = 6). **D,E.** In the same conditions, 25 pulses of light did not evoke large IPSCs from a layer 2/3 PYR in the PL (n = 10, N = 4) (**D.**) or CA1 PYR in vHPC (n = 16, N = 7) (**E.**), while in that same cell puff application of ACh (1 mM) could (insets, bottom). Heatmaps of IPSC frequency after ACh stimulation in all PL and vHPC PYRs highlight reduced sensitivity to ACh in these regions compared to the BLA.

The BLA receives extremely dense cholinergic projections from the basal forebrain, more so than other cortical structures associated with fear circuitry, notably layer 2/3 of the prelimbic cortex (PL) and the CA1 region of the ventral hippocampus (vHPC)^22,23^. While ACh is simultaneously released across these regions^17,18^ in response to a salient stimulus, the comparative effect of similar cholinergic stimulation in these circuits is not clear. Endogenous ACh can evoke rhythmic IPSCs in HPC^53,60^.

However, due to differences in cholinergic circuits, we hypothesized that large theta network activity would be less reliably evoked in the PL or vHPC compared to the BLA in response to the same cholinergic stimulus. To test this, recordings from PYRs in either layer 2/3 in the PL or CA1 of the vHPC were performed. As hypothesized, large IPSCs were much less reliably evoked in both the PL and the vHPC compared to the BLA (Figure 3D and 3E), even in the presence of physostigmine. Interestingly, in cells where light stimulation did not evoke these events, a focal puff of ACh (1 mM) was able to evoke rhythmic large events (Figure 3D and 3E), suggesting that the light stimulus failed to do so due to insufficient levels of ACh at critical points in the circuit. The findings suggest that cholinergic input preferentially induces theta network activity in BLA over hippocampus and cortex, highlighting that the same cholinergic stimulus has brain area specific effects.

### Acetylcholine activation of perisomatic inhibitory populations in the BLA

To understand how ACh modulates BLA inhibitory circuits to produce theta oscillations, we utilized the inhibitory opsin halorhodopsin (AAV-EF1a-DIO-eNpHR3.0) to selectively silence CCK, PV, or SOM IN populations during recordings of spontaneous IPSCs (sIPSCs) from BLA PYRs (Figure 4A). In baseline conditions, inhibition of SOM INs resulted in the largest reduction in total sIPSC charge and frequency compared to CCK or PV interneurons (Figure 4B), indicating SOM INs as dominant drivers of BLA inhibition in baseline states.

**Figure 4.**
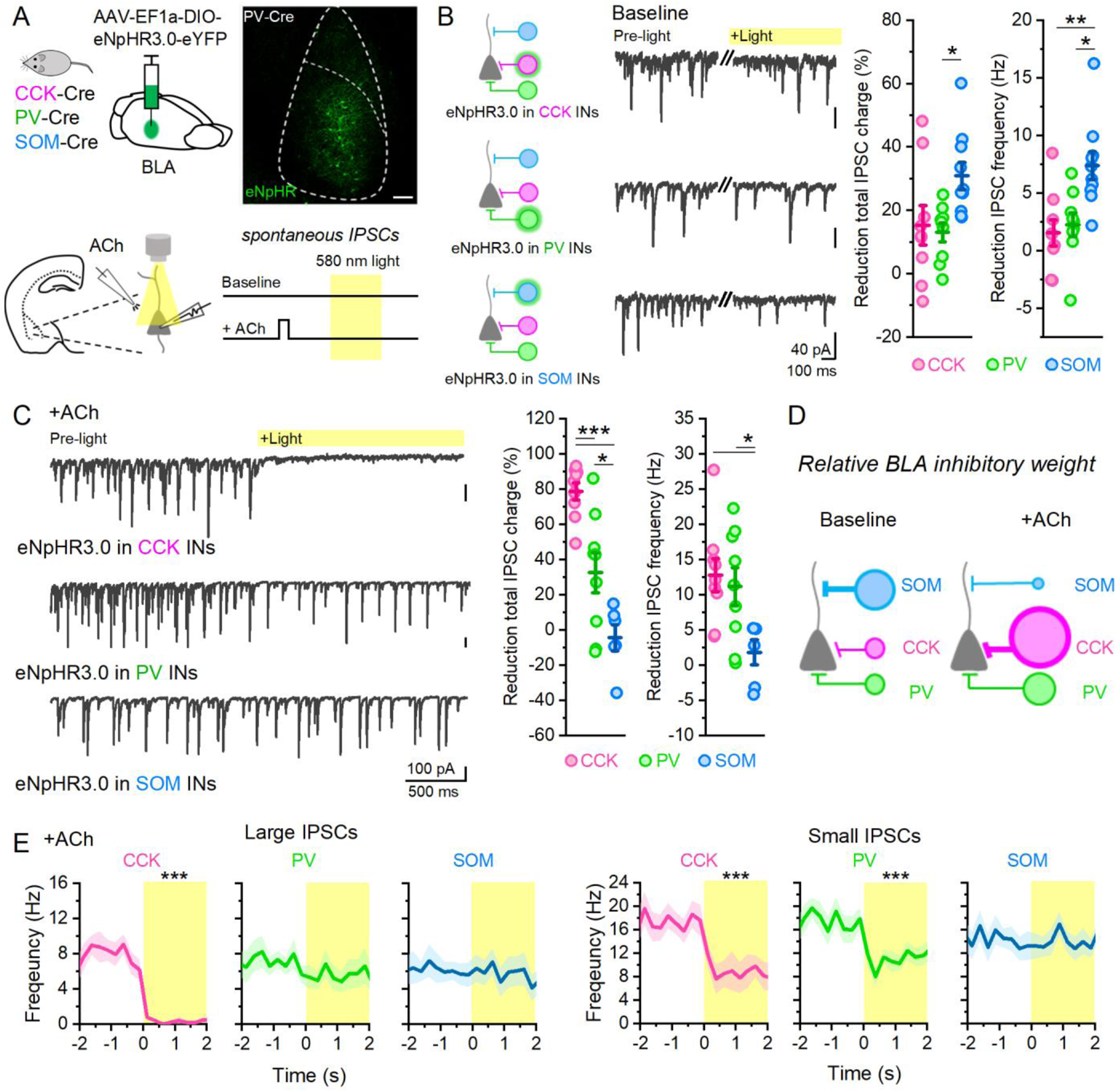
Acetylcholine shifts dominant form of inhibition in BLA. **A**. Schematic of experimental design where BLA IN populations are selectively inhibited by halorhodopsin in brain slice preparations. Scale bar 100 μm. **B**. Sample waveforms from BLA PYRs showing sIPSCs before and during light inhibition of CCK (n = 9, N = 4), PV (n = 9, N = 3), and SOM INs (n = 10, N = 4). Total reduction of IPSC charge and IPSC frequency during light is highest during SOM IN inhibition (One-way ANOVA, IPSC Charge p=0.022; IPSC frequency, p=0.002). **C**. Sample waveforms from BLA PYRs after ACh show different effects of light inhibition of CCK, PV, and SOM INs compared to baseline. Total reduction of IPSC charge and IPSC frequency during light is now highest during CCK IN inhibition (One-way ANOVA, IPSC Charge p=7.6×10^−6^, IPSC frequency, p=0.018). **D**. Schematic showing the representative shift in BLA inhibition by ACh. **E**. Comparing the frequency of large (>5x baseline average) and small (1-3x baseline average) IPSCs after ACh shows IN specific involvement.

Next, we sought to determine how ACh modulated activity in these inhibitory networks. ACh (1 mM) was briefly (500 ms) focally applied through a puffer pipette positioned near the recorded cell, which produced similar effects on BLA circuits as optogenetically released ACh (Supplemental Figure 2). Following focal ACh application, eNpHR inhibition of these different IN classes revealed a dramatic shift in activity of BLA inhibitory circuits. In these conditions, inhibition of CCK INs now blocked the large majority of IPSC charge (Figure 4C), while inhibition of PV INs blocked a smaller component.

Conversely, inhibition of SOM interneurons in this condition had no effect. There was also a larger decrease in IPSC frequency by inhibiting CCK and PV INs when compared to that of SOM INs. Taken together, these results indicate that ACh, acting via M3 mAChRs, shifts the BLA network from a state of strong SOM-mediated inhibition to a state of dominant CCK- and PV-mediated inhibition (Figure 4C, D), shifting inhibition from dendritic to perisomatic compartments.

To understand the contribution of these different IN populations to theta oscillatory activity following ACh, IPSCs were separated into large or small events and their frequency plotted over time prior to and during eNpHR inhibition (Figure 4E). Inhibition of CCK INs (n = 12, N = 4) resulted in a complete reduction of large IPSCs (paired t-test, p = 5.12 x 10^−6^) and a partial reduction of small IPSCs (paired t-test, p = 0.0015). PV IN inhibition (n = 16, N = 7) had no effect on the large IPSC frequency (paired t-test, p = 0.274) but did result in a partial reduction of the small IPSCs (paired t-test, p = 4.88 x 10^−4^). Finally, inhibition of SOM INs (n = 13, N = 6) had no effect on the frequency of either the large (paired t-test, p = 0.661) or small IPSCs (paired t-test, p = 0.671) after ACh. These results indicate that large theta frequency IPSCs induced by ACh were mediated by CCK INs, while the increased small IPSCs were produced by both PV and CCK INs. SOM INs did not contribute significantly to either type. Differences in contribution of PV and CCK INs to large IPSCs could not be explained by a different number of contacts of PV and CCK axon terminals onto BLA PYRs (Supplemental Figure 3), instead suggesting that ACh is differentially driving these perisomatic IN networks. To confirm that CCK INs also drive large IPSCs in response to optogenetically released ACh, GABA release was selectively inhibited from CCK terminals via cannabinoid receptor 1 (CB1R) activation^61^ after optogenetic stimulation of ACh (Supplemental Figure 4). These manipulations also blocked ACh-induced large IPSCs, confirming CCK INs as driving oscillatory theta inhibition in the BLA in response to large cholinergic stimulation.

### SOM interneuron activity can block CCK-induced theta inhibition in the BLA

SOM INs can fire rebound spikes following eNpHR inhibition^62,63^. Indeed, unit recordings from SOM INs in our preparations showed the presence of rebound spikes following light inhibition that could produce a barrage of rebound ISPCs in recorded PYRs in these slices (Figure 5A and B). Interestingly, while eNpHR inhibition of the SOM INs had no effect on IPSCs induced by ACh (Figure 4C and E), large IPSCs in BLA PYRs were significantly disrupted immediately by the cessation of light inhibition of SOM INs. Blockade of the large IPSCs by SOM rebound firing suggested a critical connection between SOM and CCK IN activity in the BLA, whereby SOM IN activity could block ACh-induced theta inhibition driven by CCK INs. To further test this, ChR2 was selectively expressed in SOM INs in BLA slices. This allowed for optogenetic activation of all SOM INs during ACh-induced CCK-mediated IPSCs in the BLA to determine if activation of SOM INs alone could disrupt rhythmic CCK-mediated theta inhibition (Figure 5C). Indeed, light activation of SOM INs (490 nm light, 1 second duration) after ACh blocked the large IPSCs (Figure 5D) within ms of SOM activation, suggesting that activity in these cells gate CCK IN activity in the BLA and theta oscillatory inhibition in response to ACh.

**Figure 5.**
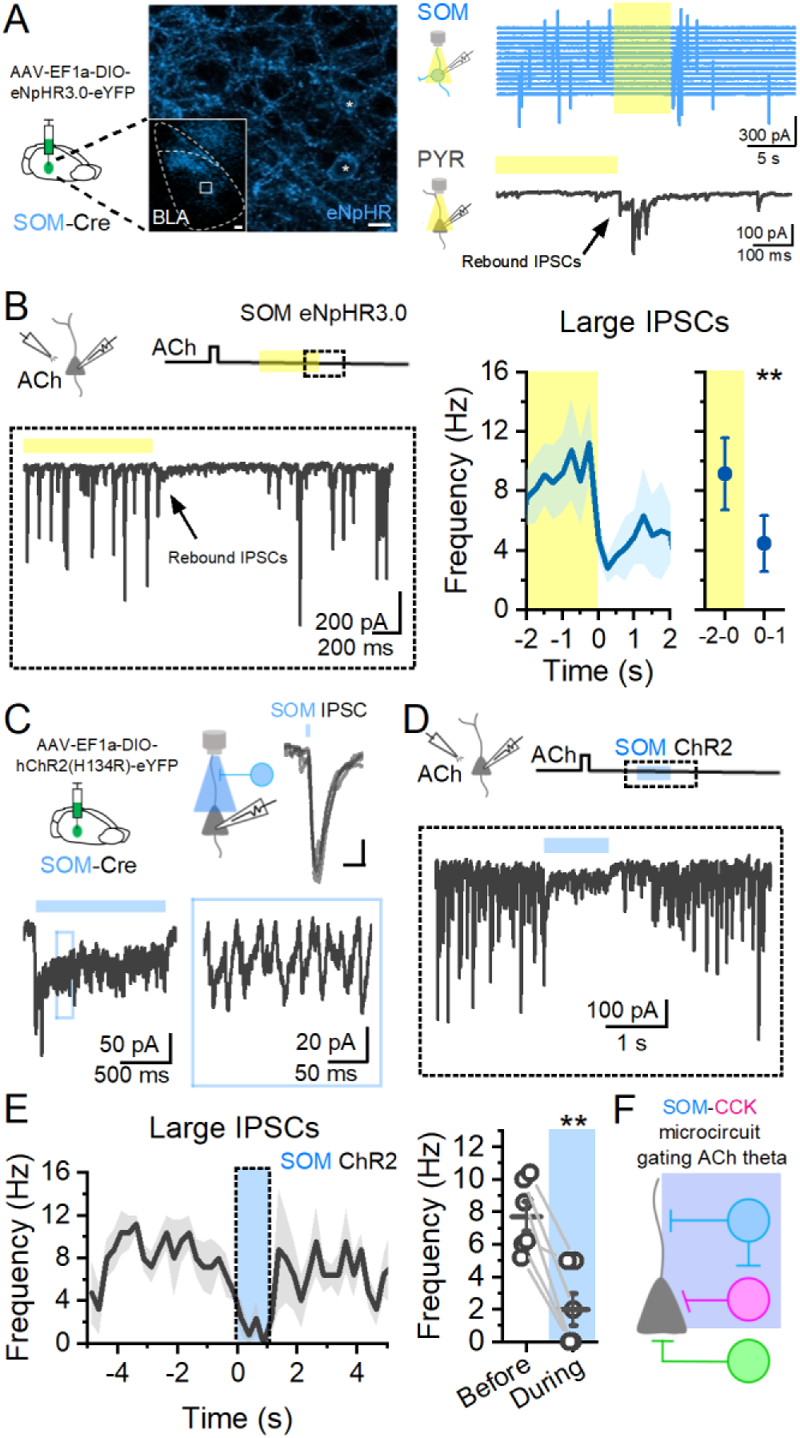
SOM INs block CCK IN-induced theta frequency inhibition after acetylcholine. **A.** Sample waveforms showing rebound firing in unit recording of SOM IN expressing eNpHR and rebound IPSCs in PYR. Scale bar 20 μm. **B**. Rebound firing of SOM INs after light inhibition produces a barrage of IPSCs in BLA PYRs that block large IPSCs after acetylcholine (n=10, N = 3; paired t-test, p = 0.004). **C**. Expression of ChR2 in SOM INs allows for optical activation of SOM INs with a single light pulse (top) or a barrage of SOM IN IPSCs during a 1 second light stimulus (bottom). **D.** Sample waveform showing activation of SOM INs disrupts CCK-mediated IPSCs in BLA PYR induced by ACh. **E**. Large IPSC frequency is decreased during activation of SOM INs (n = 6, N = 3, paired t-test, p = 0.003). **F.** Circuit schematic showing SOM innervation of CCK INs that can gate ACh-induced theta inhibition.

### Differential response to focal acetylcholine by CCK, PV, and SOM interneurons

We have shown differential activity of CCK, PV, and SOM IN networks in the BLA after ACh stimulation. To gain a mechanistic understanding of this at the cellular level, we performed fluorescently targeted whole-cell patch clamp recordings from these INs using different transgenic strategies (Figure 6A). Recordings from these preparations revealed characteristic electrophysiological differences in the cell populations (Figure 6B) consistent with previous reports^40^, confirming targeting in largely nonoverlapping populations.

**Figure 6.**
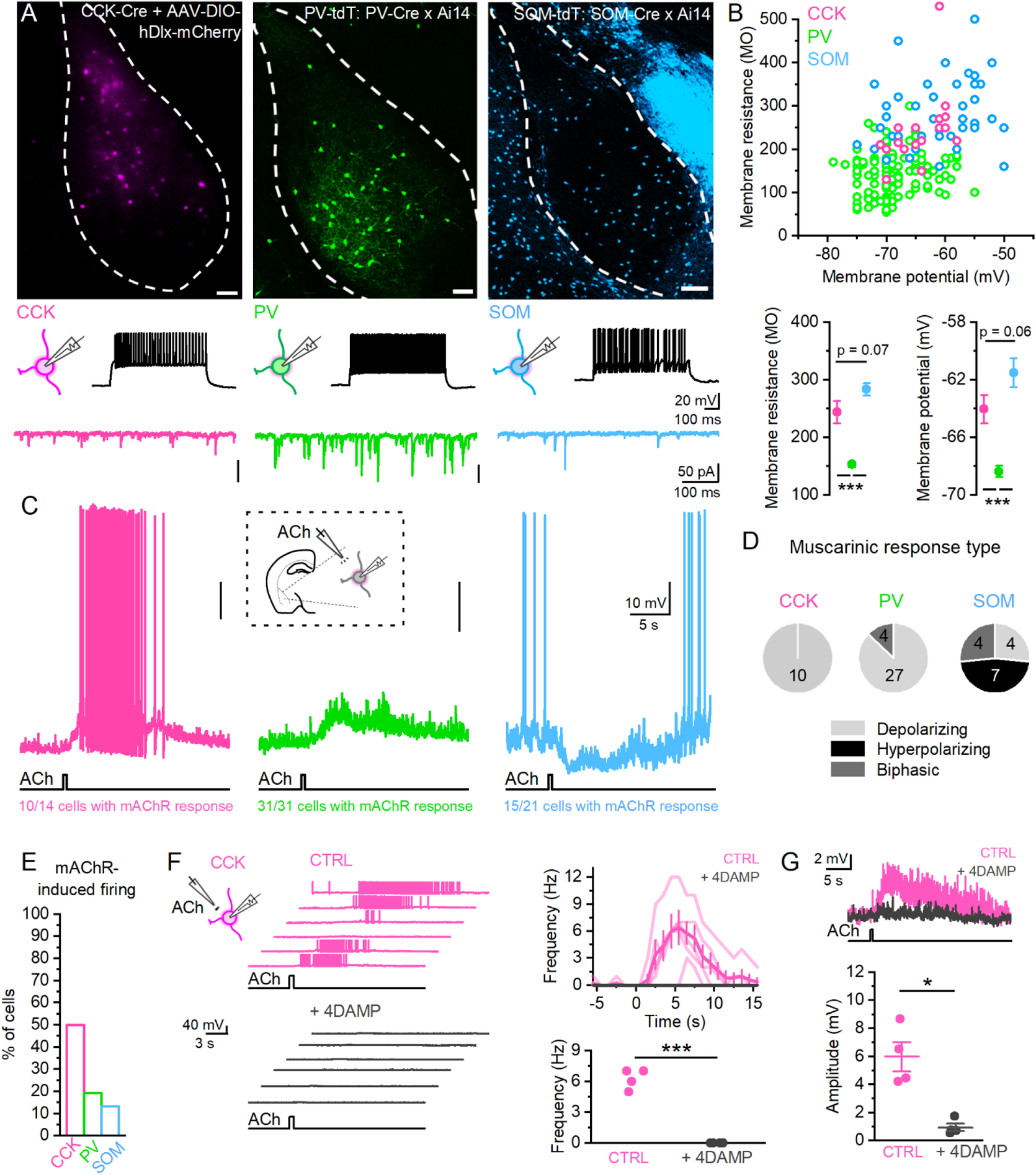
Acetylcholine differentially impacts CCK, PV, and SOM INs in the BLA. **A**. Targeted recordings from CCK, PV, and SOM INs, showing different firing patterns to current injection and spontaneous EPSCs (traces) were obtained using different transgenic strategies (top, confocal images showing LA/BLA complex, dotted white lines). Scale bars 100 μm. **B**. Recordings from tagged neurons in these preparations show that these populations of INs can be distinguished based on electrophysiological parameters. **C**. Local puff application of acetylcholine onto CCK, PV, and SOM INs reveals differential mAChR responses to ACh in these populations. **D.** Breakdown of mAChR response in each IN population. **E**. CCK INs are more likely to fire action potentials from rest in response to ACh. **F**. Sample waveform showing firing in a CCK IN in response to repeated puff application of ACh in control conditions (pink) or after application of 4DAMP (1μM). Graph (top right) showing the firing frequency in each trial (light pink) and average (dark pink) for the sample cell and the average firing frequency in all cells in control and 4DAMP (bottom right; paired t-test, p = 9.7 x 10^−4^, n = 4, N = 3). **G.** Sample waveform showing that underlying depolarization in CCK INs is blocked by 4DAMP (paired t-test, p = 0.016, n = 4, N = 3).

To explore how these IN populations are modulated by mAChRs, focal puff application of ACh (1 mM) was applied within roughly 50 microns of the recorded cell. In some cells, an initial fast nAChR response was observed. In these instances, the mAChR response was determined with mecamylamine (20μM) present. Not surprisingly, when recording from a resting membrane potential, the majority of both CCK and PV interneurons exhibited a slow depolarizing response to ACh (Figure 6C,D). For CCK INs, 71% of cells (10/14 cells) had a slow ACh response that could be completely blocked by atropine (5μM), indicating it was a mAChR response. In all CCK INs with a mAChR response, the response was depolarizing. On the other hand, while all PV INs exhibited a mAChR response (31/31 cells), a minority of the cells responded in a biphasic manner (4/31 cells), with an initial hyperpolarizing response followed by a delayed depolarization (Figure 6D). While a minority of SOM interneurons were also depolarized by ACh (4/15 cells), nearly half of all recorded SOM INs were hyperpolarized (7/15 cells). In many instances, ACh blocked spontaneous action potential firing in SOM INs. Like PV INs, a minority of SOM interneurons also responded in a biphasic manner (4/15 cells). In addition to fluorescently targeted IN recordings, recordings from putative INs (as identified by electrophysiological parameters such as firing properties and membrane resistance) in ChAT-ChR2 slices showed similar muscarinic responses to released ACh (Supplemental Figure 5). These experiments indicate that our focal puff application of ACh acts on INs in the BLA in a similar manner to transient endogenous ACh release, allowing these results to extend to endogenous ACh release.

Interestingly, from resting membrane potential, mAChR responses induced IN firing in 50% of CCK INs recorded, compared to only 19% of PV and 13% of SOM (Figure 6E). When compared to PV INs, the difference in CCK sensitivity to firing could not fully be explained by the amplitude of ACh depolarization (amplitude ACh depolarization: PV = 4.12 ± 0.49 mV, CCK = 5.97 ± 1.03 mV; two sample t-test, p = 0.09), but was also likely due to differences in intrinsic parameters in these populations, including CCK INs exhibiting a more depolarized resting membrane potential, higher input resistance, and lower rheobase to action potential firing (resting membrane potential: PV = −68.37 ± 0.41 mV, CCK = −64.05 ± 0.97 mV; two sample t-test, p = 3.16 x 10^−4^; input resistance: PV = 152.99 ± 4.66 MΩ, CCK = 243.33 ± 19.53 MΩ, p = 4.68 x 10^−9^; PV: n = 132, N = 45; CCK: n = 18, N = 7) and (rheobase: PV = 187.25 ± 19.09 pA, n = 20, N = 6; CCK = 94.70 ± 11.79 pA; n = 17, N = 7; p = 3.58 x 10^−4^). In paired PV-PYR recordings, even when PV INs did fire in response to ACh, their action potentials did not align with large IPSCs in the PYR (Supplemental Figure 6). This contrasted from large events in baseline where PV IN firing was directly aligned, further indicating a shift in the circuit mechanisms underlying synchronized network activity by ACh. Indeed, optogenetic activation of PV INs could drive theta oscillations of BLA LFP (Supplemental Figure 6) further indicating that these cells are capable of driving theta oscillations but are not doing so in response to cholinergic stimuli.

Because ACh-induced large IPSCs in BLA PYRs were mediated by CCK interneurons and were blocked by the M3 antagonist 4DAMP, we hypothesized that CCK depolarization by ACh was also mediated by M3 mAChRs. Indeed, application of 4DAMP (1uM) blocked CCK IN firing by ACh (Figure 6F) and the underlying depolarization (Figure 6G).

### Acetylcholine activates and synchronizes BLA PYRs

To understand how ACh is interacting with PYRs in the BLA during ACh-induced theta LFP oscillations, whole-cell patch clamp recordings of PYRs were performed. In the presence of glutamate and GABA receptor antagonists (20 μM CNQX, 50 μM APV, 20 μM picrotoxin, and 1μM CGP), BLA PYRs exhibited a direct biphasic response to the cholinergic stimulus: an initial hyperpolarization followed by a slow depolarization that persisted beyond light stimulation (Supplemental Figure 7), consistent with previous reports^55,56^. Interestingly, the slow depolarizing response in PYRs after light stimulation was not induced in all cells and often required increasing light stimulation or the presence of physostigmine (1 μM), again indicating that ACh’s effects on BLA circuits are dependent on the size of the cholinergic stimulus. Further, multiunit recordings show a biphasic response in unit frequency by ACh that is not dependent on glutamate or GABA signaling, but blocked by atropine (Supplemental Figure 8), indicating a direct ACh response driving changes in excitability in these cells.

**Figure 7.**
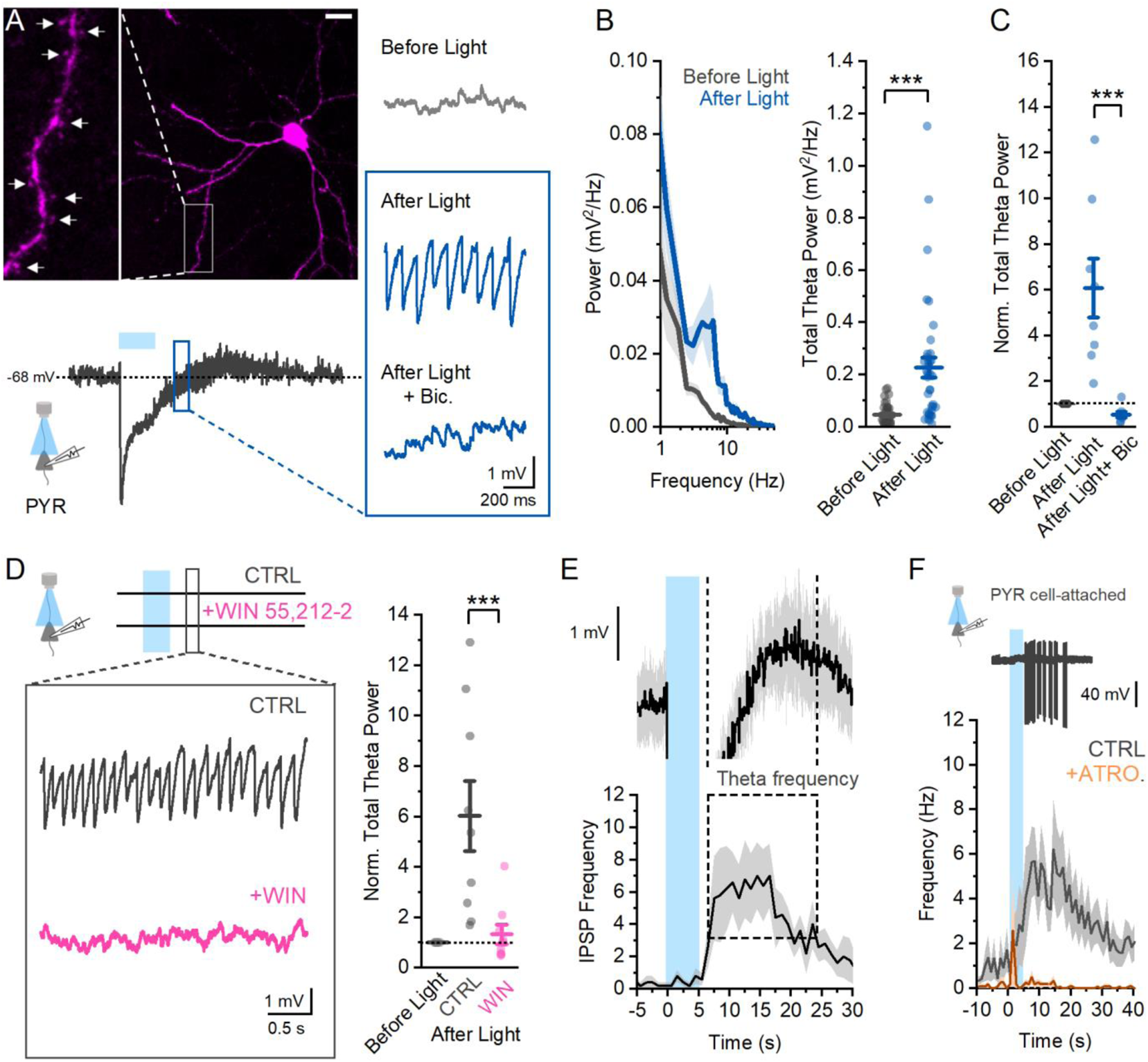
Acetylcholine produces theta frequency oscillations in BLA PYRs. **A.** An example image of a recorded PYR showing characteristic spiny dendrites (top) and biphasic post-synaptic response to released ACh. Magnification of the membrane potential shows large rhythmic activity after light that is blocked by bicuculine (20 μM). Scale bar 25 μm. **B.** Average power spectrums of PYR membrane potential before and after light show large increase in theta power (3-12 Hz) after light (paired t-test, p = 2.28 × 10^−5^, n = 38, N = 13). **C.** This increase in theta power is blocked by bicuculline (n=8, N=5, one-way repeated measures ANOVA, p = 5.24×10^−4^). **D.** Application of the CB1 receptor agonist WIN 55,212-2 (2uM) also blocks theta oscillation in BLA PYRs induced by ACh (n=9, N=3, one-way repeated measures ANOVA, p = 0.032). **E.** Plotting the average BLA PYR membrane potential after ACh stimulation shows a temporal overlap with theta frequency IPSPs in these cells. **F.** Unit recordings from BLA PYRs show that in response to cholinergic stimulation they can fire at a theta frequency, which is blocked by atropine (5 μM) (average unit frequency after light = 4.66 ± 0.98 Hz; n = 11, N = 7; peak frequency after light + atropine = 0.21 ± 0.14 Hz; n = 4, N = 3, two sample t test, p = 0.022.)

With local glutamate and GABA transmission intact, in addition to the direct cholinergic response, BLA PYRs also exhibited a slow-onset large theta frequency membrane potential oscillation (MPO) after light stimulation (Figure 7A, B). In agreement with IPSC experiments, application of the GABA_A_ receptor antagonist bicuculline (20 μM) completely blocked the theta MPOs induced by light (Figure 7C). Consistent with a role of CCK INs driving theta inhibition in this state, MPOs in PYRs were also abolished by the CB1 receptor agonist WIN 55,252-2 (2 μM) (Figure 7D).

Interestingly, we noted that large IPSPs evoked by ACh reached theta frequency during the rising phase of the ACh-induced depolarization of BLA PYRs (Figure 7E). To test the hypothesis that this temporal relationship would have implications for spiking output of BLA PYRs, we performed cell-attached unit recordings in BLA PYRs (Figure 7F). As predicted, ACh caused increased action potential firing that peaked in the theta frequency range and could be blocked by atropine (average unit frequency after light = 4.66 ± 0.98 Hz; n = 11, N = 7; peak frequency after light + atropine = 0.21 ± 0.14 Hz; n = 4, N = 3, two sample t test, p = 0.022.

Coactivity in ensembles of BLA PYRs is important in activating down-stream structures and mediating amygdalar behaviors. Theta oscillations are a mechanism by which local activity is synchronized in a circuit. To explore how ACh-induced theta oscillations modulate BLA PYR synchrony, paired recordings of neighboring (within < 100 microns) PYRs were made (Figure 8A). In baseline conditions, neighboring cells exhibited low correlation between fluctuations in their membrane potential (Figure 8B). After ACh stimulation, theta MPOs driven by large IPSPs greatly synchronized these cells (Figure 8B). Across 23 pairs of BLA PYRs, 15 pairs (65%) exhibited high (> 0.25) cross correlation of ACh-induced MPOs. The mechanism underlying MPO synchronization in PYR pairs could be explained by large theta IPSCs (Figure 8C). In a different set of recorded pairs, large theta IPSCs were found to be synchronized across 12 of 19 pairs (63%). In the remaining 7 pairs, large IPSCs induced by ACh were present in each cell but not synchronized (7/19, 37%). These results suggest that ACh synchronizes the majority of neighboring BLA PYRs via shared connectivity within local CCK IN circuits, but leaves the possibility that there are some PYRs that are differently connected within the local inhibitory circuitry.

**Figure 8.**
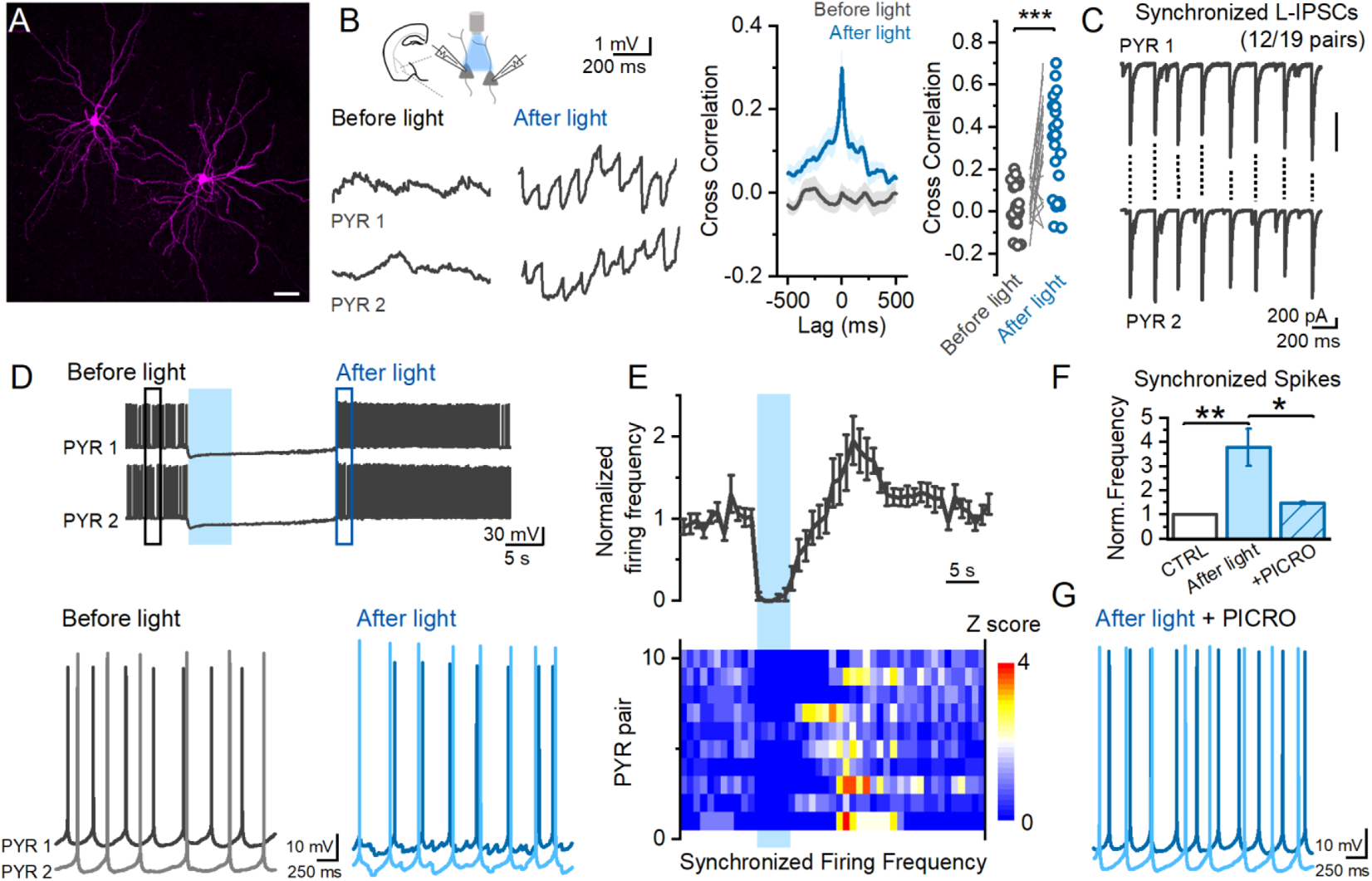
Acetylcholine synchronizes BLA PYRs. **A**. Confocal image showing an example of a dual neighboring PYR recording in the BLA. Scale bar 50 μm. **B.** Representative paired recordings of PYRs showing the high cross correlation of membrane potentials during ACh-induced MPOs that occurs in most, but not all pairs (Right, paired t-test, p = 6.25 x 10^−5^, n = 23, N = 12). **C.** In voltage clamp recordings, synchronization of ACh-induced large IPSCs can be observed in most cells, providing a mechanism for synchronized MPOs. **D,E.** Sample recordings showing that when neighboring PYRs are brought to firing threshold, acetylcholine induces an inhibition of firing activity that is followed by an increase in the synchronization of PYR action potentials. **F**. Synchronization of spiking induced by acetylcholine (n = 10 pairs, N = 8) is blocked in the presence of picrotoxin (n = 3 pairs. N = 3) _to block_ GABAA receptors (two sample t-test, p = 0.01), shown in **(G)** traces from the same cell pair.

Finally, to address how synchronized MPOs across PYRs could modulate spiking synchronization, paired recordings of neighboring PYRs were given current injections to induce sustained low frequency firing (range: 2-5 Hz). In baseline conditions, the frequency of synchronized spikes (spikes less than 20 ms apart) was low (Figure 8D). ACh stimulation evoked an initial inhibition of firing followed by firing at an increased frequency from baseline (Figure 8E) in which spiking synchrony across PYR pairs was significantly increased (Figure 8F). This increase in firing synchrony was blocked by application of picrotoxin to block GABA_A_ receptors (Figure 8F, G), indicating that the increase in firing synchrony was produced by ACh-induced CCK theta inhibition and could not be explained by an overall increase in spiking output, as firing frequency after light + picrotoxin was still increased from baseline.

## Discussion

Acetylcholine plays a vital role in establishing network states in the brain that support attention, memory, and emotional behaviors^6–10^. These high ACh states are associated with theta oscillations which bind activity of neuronal ensembles within and between brain regions. In the BLA theta oscillations are critical for emotional behaviors^24–28^. However, despite the remarkably dense cholinergic innervation of the BLA, a mechanistic understanding of how ACh acts in the BLA to produce theta oscillations is lacking. Here, our data collectively outline a detailed temporal and cell-specific mechanism by which endogenous phasic ACh release induces theta oscillations in the BLA through a previously undefined microcircuit mechanism. Additionally, we show that endogenous ACh stimulation alters BLA circuits in a stimulus-sensitive manner and that the same stimulation more readily modulates BLA compared to vHPC or PL circuits.

### SOM INs gate CCK-driven theta oscillations in BLA

While PV INs have been implicated in BLA theta oscillations^48–50^, it is not clear if other IN populations can also play a role. Similar to PV INs, CCK INs heavily innervate the perisomatic region of BLA PYRs and can modulate fear behavior^64^. However, the role of CCK INs in network function has remained elusive. Utilizing multiple approaches, including targeted optogenetic silencing, pharmacology, and targeted whole cell recordings, we show that CCK INs are recruited by cholinergic stimuli to synchronize PYR output and drive theta oscillations in the BLA, establishing a new role for these cells in BLA circuit function. Large CCK-mediated IPSCs induced by ACh contained tightly summated IPSCs, suggesting phasic cholinergic stimulation results in highly synchronized activity in the CCK IN network. This synchronized inhibition was sensitive to CB1 receptor manipulation, indicating it is likely driven specifically by the CCK basket cell network^65^, as there exist multiple classes of CCK INs in the BLA^66^. Rhythmic inhibition in HPC induced by ACh is also sensitive to CB1Rs^53,60^, suggesting this cell type could play a similar role in other brain areas. CCK basket cells in the BLA are electrically coupled^44^, providing a mechanism by which M3 receptor depolarization can drive synchronized output from this network without the need for glutamatergic signaling. An individual CCK IN is estimated to innervate 700-800 local PYRs^67^, enabling synchronization of this CCK IN network to entrain a large ensemble of BLA PYRs and modulate BLA LFP activity. Indeed, the majority of paired PYR recordings showed synchronized inhibition that entrained spiking output. However, not all neighboring PYRs were synchronized in this state, suggesting that ACh-induced theta oscillations may differentially modulate BLA PYR networks. Intriguingly, specific projection pathways are differentially regulated by CCK INs in the BLA^68^, indicating that ACh-induced theta oscillations could separate BLA projection outputs. Future studies with projection-specific recordings will be needed to examine this possibility.

In the BLA, SOM INs have been implicated in modulating external inputs^45^, plasticity^69^, and feedforward^70^ and feedback inhibition^71^. Here, we describe a novel role of SOM INs in gating activity of CCK INs to modulate BLA network synchronization. SOM INs exhibit high levels of activity in baseline states in cortical circuits *in vivo*^72,73^ and we show this extended to BLA *ex vivo*. ACh decreased activity in this IN population, directly hyperpolarizing a large subset of these cells. This contrasts with the effects of ACh on SOM INs in other brain areas^74^ but is consistent with decreased BLA SOM activity following systemic physostigmine^46^. This ACh-induced reduction in SOM activity shifts the network state in the BLA from high dendritic to strong perisomatic inhibition and allows for a CCK-driven theta oscillation to emerge. Interestingly, our data show that activation of SOM INs in this network state blocks CCK IN oscillatory inhibition, establishing a previously undefined functional relationship between SOM and CCK INs. They also establish that this SOM-CCK IN circuit must be suppressed to allow for CCK-mediated synchronization of BLA PYR theta activity. This provides an important role for ACh in inhibiting SOM INs to allow this network behavior and positions these INs to gate theta oscillations in the BLA. Intriguingly, ACh-mediated inhibition of SOM INs not only gates theta oscillations, but also disinhibits PYR dendrites, supporting plasticity on PYR cell dendrites during BLA theta activity. Inhibition of SOM INs is required for plasticity of external inputs to BLA PYRs^69^. By inhibiting SOM INs, ACh may therefore enable both theta oscillations and plasticity in BLA. Indeed, ACh plays an important role in emotional learning^8,10^.

### ACh shifts control of BLA theta oscillations from PV to CCK INs

PV INs in the BLA are capable of driving theta oscillations^48–50^. However, we show that despite PV IN activity being increased by ACh, these INs were not involved in the theta network behavior produced by phasic cholinergic stimulation, despite their involvement in network activity in baseline states. This suggests their involvement in network oscillations in the BLA is state-dependent. PV and CCK IN networks in the BLA form parallel circuits without direct synaptic communication^44^, unlike the hippocampus^75^. While PV and CCK cells have differences in electrophysiological parameters, they equally inhibit BLA PYR firing^67^. In any brain area, there is limited exploration on how complementary PV and CCK IN networks, both innervating the perisomatic region, differentially contribute to circuit dynamics^76^. What is the benefit of these two competing populations in modulating circuit activity? One explanation is that these networks are differentially recruited to circuit function and play a state-dependent role. PV INs receive more glutamatergic inputs than CCK INs, while CCK INs are more sensitive to neuromodulatory control^40,77,78^. Accordingly, external glutamatergic inputs, such as those from cortex, could drive synchronization in BLA through interaction with PV INs. Alternatively, ACh suppresses cortical glutamatergic input^52^ and promotes a network state in BLA in which CCK INs internally drive theta oscillations without the need for external glutamatergic input. This suggests externally and internally driven theta oscillations in the BLA could be generated by these competing IN networks. Consistent with this, our data show that due to differences in intrinsic parameters in CCK INs, these cells are more sensitive to ACh stimuli than PV INs in the BLA. Thus, in response to phasic cholinergic stimuli, CCK INs provide stronger control over the perisomatic domain and synchronize BLA output.

In the presence of ACh, increases in PV IN activity did not align with large CCK-mediated IPSCs, highlighting the independent nature of these two circuits in this state. PV INs are key players in gamma oscillations in BLA circuits through interactions with local PYRs^79^. Increased excitation of PV INs by ACh, in combination with ACh-evoked depolarization of PYRs, could generate gamma oscillations in the BLA. In agreement, gamma power was increased in BLA by ACh and while this was not examined in detail in the current study, our results suggest that PV INs could play a role. Interestingly, this could offer parallel, separate control of theta and gamma oscillations in the BLA through CCK and PV networks by phasic ACh.

### Acetylcholine effects on circuit behavior are cholinergic stimulus- and brain area-specific

Basal forebrain cholinergic neurons respond to emotionally salient stimuli, resulting in phasic release of ACh into the BLA^15^ simultaneously with hippocampal and cortical areas^17,18^. Firing in cholinergic neurons can vary based on the value of salient stimuli^17^ and different ACh levels have been hypothesized to set network states for different functions^80^. The BLA receives especially dense cholinergic innervation compared to these other structures^22,23^. Here, our data show that induction of theta oscillatory network behavior requires large cholinergic stimuli and due to this, the same stimulation to cholinergic fibers more reliably recruits the BLA compared to vHPC, and PL. This indicates a threshold of ACh must be reached to induce this network behavior. These results have intriguing implications. First, depending on the level of innervation by cholinergic fibers, this results in differential effects in target structures by the same cholinergic stimulus, offering important insight into the behavioral effects of ACh signaling. In response to salient stimuli, this positions ACh to more readily induce synchronized theta behavior in the BLA important for the transmission of valence information. Indeed, due to strong cholinergic innervation of BLA our work shows that it is possible for a single light pulse, resulting in synchronous activation of cholinergic fibers as is observed in BF in response to a salient stimulus^16^, to induce this network behavior. Interestingly, cholinergic stimulation first results in a decrease in BLA network activity. It is possible this serves to reset BLA activity prior to induction of theta oscillations that are generated locally in BLA circuits without external glutamatergic signaling. Indeed, BLA theta oscillations can lead synchronized behavior in vHPC and PL circuits during emotional processing^33^. It is an intriguing possibility that this is driven by differential cholinergic modulation in these regions. Additionally, stimulus specificity of ACh’s effects on BLA network oscillations could allow only salient stimuli to induce this network state as opposed to weak stimuli that lack strong valence.

## Conclusion

In conclusion, our findings outline the cell-specific circuit mechanism by which endogenous phasic ACh modulates BLA theta oscillations and synchronizes BLA output acting through CCK INs. These results show how ACh modifies inhibitory circuits in the BLA to establish a network state favoring theta oscillations and reveal a previously undefined relationship between CCK and SOM INs that is critical for modulating oscillatory activity. We also show that ACh preferentially impacts BLA circuits over other hippocampal or cortical circuits in this manner. While not the only mechanism by which BLA circuits produce theta oscillations, this likely represents an important mechanism by which BLA theta oscillations are produced in response to presentation of salient environmental stimuli that plays an important role in emotional learning and memory.

## Acknowledgements

We thank all members of the Mott laboratory for their comments on this project. This work was supported by National Institute of Mental Health R01MH104638 to D.D.M. and A.J.M.; University of South Carolina VP for Research ASPIRE2 Award to D.D.M.; University of South Carolina VP for Research Support to Promote the Advancement of Research and Creativity (SPARC) Research Grant to J.X.B-P.

## Contributions

J.X.B-P and D.D.M designed research. J.X.B-P collected electrophysiology data, J.X.B-P and G.C.J collected anatomical data. J.W.W performed viral injection surgeries. J.X.B-P and D.D.M analyzed data. J.X.B-P and D.D.M wrote the paper. J.X.B-P, D.D.M, and A.J.M edited the paper. J.X.B-P, D.D.M, and A.J.M obtained funding for the research.

## Declaration of Interest

The authors declare no competing interests.

## METHODS

### Animals

All animal use and care procedures were performed under the guidelines of the National Institute of Health’s *Guide for the Care and Use of Laboratory Animals* (Department of Health and Human Services) and approved by the University of South Carolina Institutional Animal Care and Use Committee. Mice had access to *ad libitum* food and water and were socially housed on a 12hr light/dark cycle. Multiple transgenic mouse lines were purchased and bred for use in this study. For experiments utilizing optogenetically release acetylcholine, female Chat-Cre mice (B6;129S6-Chat ^tm2(cre)Lowl^/J, stock number #006410, The Jackson Laboratory) were crossed with male Ai32 mice (B6; 129S-Gt(ROSA)^26SOR^ ^tm21(CAG-COP4*H134R/EYFP)Hze^ /J, #012569, The Jackson Laboratory) to create ChAT-ChR2 offspring^55,56^. For targeted interneuron recordings and manipulations the following transgenic mice were used: PV-tdTomato mice created by crossing female PV-Cre (B6;129P2-*Pvalb^tm1(cre)Arbr^*/J,#008069, The Jackson Laboratory) with male Ai14 (B6.Cg-*Gt(ROSA)26Sor^tm14(CAG-tdTomato)Hze^*/J,#007914, The Jackson Laboratory), SOM-tdTomato mice created by crossing female SOM-Cre (*Sst^tm2.1(cre)Zjh^*/J,#013044, The Jackson Laboratory) with male Ai14, and CCK-Cre mice (C57BL/6-Tg(Cck-cre)CKres/J,#011086, The Jackson Laboratory). Both male and female mice of all transgenic lines were used in this study when they were between 5 – 24 weeks old.

### Stereotaxic viral injections

For stereotaxic surgeries, mice that were at least 6 weeks old were anesthetized with isoflurane and secured in a stereotaxic apparatus (Stoelting). Adeno-associated virus (AAV, 150 – 350 nL) was slowly injected into the BLA of both hemispheres (coordinates relative to bregma: anterior/posterior: −0.4 mm, medial/lateral: ± 3.2 mm, dorsal/ventral: −5.2 mm). For halorhodopsin experiments, PV-tdT and PV-Cre, SOM-tdT, or CCK-Cre mice were injected with AAV5-EF1a-DIO-eNpHR3.0-eYFP (UNC Vector Core). For channelrhodopsin experiments, CCK-Cre, PV-Cre, or SST-Cre mice were injected with AAV5-EF1a-DIO-hChR2(H134R)-eYFP (UNC Vector Core). For selective florescent expression in CCK interneurons, CCK-Cre mice were injected with AAV5-hDlx-Flex-dTomato-Fishell_7 (Addgene)^81^, which selectively labels CCK interneurons. After viral injection, skin incisions were closed with topical GLUture adhesive (Abbott Laboratories) and mice were closely monitored for recovery. Injected animals were given at least 3 weeks for viral expression after surgery before use in brain slice experiments.

### Brain slice electrophysiology

For brain slice preparation, mice were deeply anesthetized with isoflurane and the brain swiftly removed and submerged in ice-cold “cutting” ACSF containing (in mM): 110 choline chloride, 2.5 KCl, 25 NaHCO_3_, 1.25 NaH_2_PO_4_, 20 glucose, 5 MgSO_4_, and 0.5 CaCl_2_. The cutting solution was continually bubbled with 95% O_2_ and 5% CO_2_. After at least one minute of recovery in this cutting solution, coronal slices of either 300 microns (whole-cell and single unit recordings) or 400-500 microns (LFP and multi-unit recordings) were made using a VT1000S vibratome (Leica). After cutting, brain slices were bisected and transferred to an incubation chamber filled with “incubating” artificial cerebral spinal fluid (ACSF) containing (in mM): 125 NaCl, 2.7 KCl, 25 NaHCO_3_, 1.25 NaH_2_PO_4_, 10 glucose, 5 MgSO_4_, and 1 CaCl_2_ superfused with 95% O_2_ and 5% CO_2_ at 34°C (pH 7.3; 300-310 mOsm). After 30 – 45 minutes of incubation at 34°C for slice recovery, the incubation chamber containing the brain slices was allowed to equilibrate to room temperature for at least 15 minutes before recordings were made.

For electrophysiological recordings, slices were transferred to a recording chamber where they were continuously perfused at a rate of 2-4 mL/minute with warm, oxygenated recording ACSF (32 - 34°C, strongly bubbled with 95% O_2_/5% CO_2_) containing (in mM): 125 NaCl, 2.7 KCl, 1.25 NaH_2_PO_4_, 25 NaHCO_3_, 10 glucose, 2 CaCl_2_ and 1 MgSO_4_ (pH 7.3; 300-310 mOsm). For whole cell recordings, neurons were visualized through a 40X water-immersion objective lens with infrared-differential interference contrast optics (Olympus BX51WI). For targeted interneuron recordings, tdTomato-expressing cells were first identified with LED excitation (pE-4000, CoolLED) and a florescent filter cube (ThorLabs). Whole-cell patch clamp recordings were performed using borosilicate glass electrodes with an input resistance of 3-6 MΩ. For LFP and multi-unit recordings, the tips of the same borosilicate glass electrodes were manually broken off to yield an input resistance of 1-2 MΩ. Input and series resistance were monitored throughout recordings and cells discarded if either changed significantly during the recording. For optogenetic experiments, LED light (pE-4000, CoolLED) of either 490 nm (ChR2 activation) or 580 nm (halorhodopsin inhibition) were applied through the 40X objective over the recorded cell. 490 nm light pulses were 1-2 ms in duration and given in 5 Hz stimulation patterns for release of acetylcholine from ChR2+ axons. For optogenetic activation experiments, ChR2 stimulation was applied with at least a 90s delay between stimulation trials for full recovery of cholinergic fibers. For optogenetic inhibition experiments, constant 580 nm light was similarly applied over the recorded cell for 2-7 seconds with at least 60 seconds between epochs of light delivery. All recordings were made with a Mulitclamp 700A amplifier (Molecular Devices) filtered at 2 kHz and digitized with a Digidata 1440A A-D board (Molecular Devices). LFP and multiunit recordings were amplified by 1000-2000x. pClamp 10 software (Molecular Devices) was used for visualization and analysis of recorded traces.

#### LFP and multi-unit recordings

The BLA is non-linear in its organization and exhibits low activity in brain slices, making LFP recordings challenging to obtain and very small in amplitude. To increase BLA slice activity in baseline, LFP and multiunit recordings were carried out in a recording ACSF solution containing elevated KCl (3.3 mM as opposed to 2.7 mM) and an increased perfusion rate (7-12 mL/minute as opposed to 2-4 mL/minute)^82^. Recording electrodes were filled with the same recording ACSF as perfused over the slice. Continuous recordings were made with optogenetic cholinergic stimulation given at 5 Hz for 5 seconds (490 nm light, 1-2 ms light pulse duration) every 90 seconds. LFP recordings were high pass filtered above 1 Hz for power spectrum analysis (pClamp 10) done in 2 second time bins directly before, during, and within 5 s after light stimulation. Total theta power was calculated as the sum of all power values from 3-12 Hz while total gamma power was calculated as the sum of all values from 30-70 Hz. Multiunit recordings were high pass filtered at 50 Hz and units detected using threshold event detection (pClamp 10). Multiunit frequency was determined as the number of units in 1 second time bins during recordings.

#### Voltage clamp post-synaptic current recordings

Voltage clamp recordings of PYRs were made using a symmetrical-chloride internal solution containing (in mM): 140 CsCl, 10 HEPES, 3 QX-314, 1 BAPTA, 4 MgATP, 0.3 Na2GTP, yielding a chloride reversal potential of 0 mV. IPSCs were isolated by application of D-2-amino-5-phosphopentanoic acid (D-APV, 50 μM, HelloBio, Princeton, NJ, USA) and 6-cyano-7-nitroquinoxaline-2,3-dione disodium salt (CNQX, 20 μM, HelloBio, Princeton, NJ, USA) to block NMDA and AMPA receptor-mediated EPSCs. Application of GABA_A_ receptor antagonists bicuculline (Bic., 20 μM, HelloBio, Princeton, NJ, USA) or picrotoxin (50 μM, HelloBio, Princeton, NJ, USA) were used to block IPSCs to confirm they were GABAergic in nature. Template event detection (pClamp 10) was used for IPSC frequency and kinetic analysis. Total IPSC charge was determined as the sum of the area of all IPSCs. Small and large IPSCs were designated relative to the average IPSC amplitude of at least 20 seconds of baseline activity, as determined using event detection. Small IPSCs were classified as IPSCs within 1-3x average baseline IPSC amplitude and large IPSCs designated as IPSCs greater than 5x the amplitude of baseline average IPSCs. For baseline halorhodopsin experiments to determine the contribution of different IN populations to sIPSCs, 580 nm light was applied multiple times for 2-7 seconds in duration for a total at least 12 seconds over the course 3-10 minutes to determine effects on IPSCs. Light on epochs were compared to IPSC activity directly preceding light and averaged over multiple trials for each cell. For acetylcholine halorhodopsin experiments to determine the contribution of different IN populations to sIPSCs, light was applied for at least 2 seconds during periods of increased IPSC activity induced by a 500 ms puff of acetylcholine (1 μM, Millipore Sigma, St. Louis, MO) through a picospritzer. Light application was repeated over multiple acetylcholine trials and averaged for each cell. Slices were only used for halorhodopsin experiments if eNpHR3.0-EGFP expressed strongly in the BLA around the recorded cell or if the recorded PYR received rebound IPSCs upon cessation of light, indicating functional halorhodopsin expression in the slice. For experiments using ChR2 in SOM INs, a pulse of 1 second 490 nm light was applied during a period of large IPSCs following focal ACh application.

#### Current clamp recordings

For interneuron recordings an internal solution containing (in mM) 135 K-gluconate, 5 KCl, 10 HEPES, 2 MgCl2, 2 MgATP, 0.3 NaGTP, and 0.5 EGTA (pH 7.3, 290 mOsm) was used. Cell parameters and spiking characteristics (in response to current injections) were determined immediately after breaking into the cell. For determining acetylcholine response, cells were held at their resting membrane potential and focal application of acetylcholine (500 ms, 1 mM in a recording ACSF solution) delivered within 50 microns from the cell body (for tdTomato identified interneuron recordings) or light stimulation of cholinergic fibers at 5 Hz for 5 seconds (for blind putative interneuron recordings in ChAT-ChR2 mice). For pyramidal neuron recordings an internal solution containing (in mM) 137 K-gluconate, 3 KCl, 10 HEPES, 2 MgCl2, 2 MgATP, 0.3 NaGTP, and 0.5 EGTA (pH 7.3, 290 mOsm) was used for detection of GABA_A_-mediated IPSPs (chloride reversal at −80 mV). PYR responses to acetylcholine were determined at a −65mV membrane potential. For experiments examining theta frequency membrane potential oscillations in PYRs, cells were held between –65mV and –55mV. For IPSP detection, pClamp template event detection was used and total IPSP frequency determined in 1 second time bins prior to and after light stimulation.

#### Paired PYR recordings

Paired recordings of neighboring PYRs (within 100 microns) were made to look at synchronization by acetylcholine. For current clamp recordings, synchronization of membrane potential before and after acetylcholine stimulation was done through cross correlation analysis (pClamp 10) in 2 second time bins. Cross correlation was measured within 20 s before and 10 s after acetylcholine stimulation in each cell pair and averaged across at least three cholinergic stimulation trials to produce a single value in that pair. This was done for both current clamp membrane potential synchronization as well as voltage clamp IPSC synchronization. PYR pairs were considered not synchronized by acetylcholine if their cross-correlation value was less than 0.25. For spiking synchrony experiments, current was injected into each recorded cell to bring that cell to spike threshold and a sustained firing rate of 2-5 Hz. Spike timing was determined by event threshold detection (pClamp) and designated as synchronized if the action potential across cells was separated by less than 20 ms. Frequency of synchronized spiking was normalized to baseline conditions and compared to the frequency of spiking synchrony after acetylcholine for each cell pair.

For all experiments, drugs were bath applied in the recording ACSF and allowed to wash on the slice for at least 2 minutes to achieve a steady state effect. In all experiments involving acetylcholine, a small amount of physostigmine (1 uM, HelloBio, Princeton, New Jersey) was present in the recording solution unless otherwise noted (Figure 3).

### Imaging and Immunofluorescence

To validate ChR2 expression in basal forebrain cholinergic neurons, mice at least 8 weeks old were transcardially perfused with ice-cold PBS containing 0.5% nitrite followed by 4% PFA. Brains were postfixed overnight in 4% PFA at 4°C before 50μm coronal brain sections were cut using a vibratome (VT1200S, Leica). Slices were blocked (TBS with 0.5% Triton X-100 and 10% donkey serum) for 30 minutes and then incubated for 48 hours at room temperature in goat anti-ChAT primary antibody (1:1000, AB144P, Millipore). Following rinse, sections were incubated at room temperature for 3 h in TBS containing AlexaFluor-546-conjugated donkey anti-goat IgG secondary antibody (1:400, A-11056, Fisher Scientific), 0.5% Triton X-100, and 2% normal donkey serum. Sections were rinsed and mounted on slides with ProLong diamond antifade mountant (Fisher Scientific). Images were captured on a Leica SP8 Multiphoton confocal microscope (Leica Microsystems) and analyzed using ImageJ (NIH) software with manual counting of antibody-fluorescence overlap.

CCK INs were fluorescently labelled by a Cre-dependent viral injection in CCK-Cre mice (AAV-hDlx-Flex-tdTomato)^81^. For targeted PV and SOM recordings, we utilized a double transgenic cross to create PV- and SOM-tdTomato mice. PV- and SOM-tdT mice were validated in a similar manner to ChAT-ChR2 mice, instead using rabbit anti-parvalbumin primary antibody (1:15,000, PV27, Swant) or rat anti-somatostatin primary antibody (1:500, sc-47706, Santa Cruz).

For experiments using *post-hoc* cell visualization, 0.1-0.3% biocytin was included in the recording pipette internal solution during whole cell recording. Brain slices were immediately transferred to a fixative solution (4% paraformaldehyde (PFA) in phosphate buffer saline) at the completion of cell recording and allowed to fix overnight. Slices were subsectioned to 50 μm using a vibratome (VT1200, Leica) and Alexa Fluor 488 Streptavidin (ThermoFisher) applied to label the recorded cell. Images were captured on a Leica SP8 multiphoton confocal microscope.

## Data Analysis

pClamp 10 and OriginPro 2018 and 2020 software were used to analyze electrophysiology data. OriginPro 2020 was used to create waveforms and graphs from the data. Data are shown as mean ± SEM. Statistical significance was determined using a t test (paired or unpaired), a one-way ANOVA, or a repeated-measures ANOVA with appropriate post hoc tests (α < 0.05 was taken as significant). Statistical significance is designated as * p < 0.05, ** p < 0.01, and *** p < 0.001. For each experiment the number of animals (N) and cells/slices (n) values are reported. Box and whisker plots with individual data points (open circles) show 25th and 75th percentiles, median (solid line) and mean (solid square).

## SUPPLEMENTAL INFORMATION

**Supplemental Table 1.** Statistical tests and expanded results.

| Figure | Parameter | Sample Size (n cells, N animals) | Primary Statistical Test | Post-hoc Test | Comparison | P | F/t statistic |
| --- | --- | --- | --- | --- | --- | --- | --- |
| 1D | Total Theta Power | 10,6 | Paired t test |  | During light vs Baseline | 0.039* | t(9) = 2.40 |
|  |  |  | Paired t test |  | After light vs Baseline | 0.006*** | t(9) = -3.56 |
| 1D | Total Gamma Power | 10,6 | Paired t test |  | During light vs Baseline | 0.77 | t(9) = 0.29 |
|  |  |  | Paired t test |  | After light vs Baseline | 0.037* | t(9) = -2.44 |
| 2C | IPSCs - Total Theta Power | 12,5 (CNQX/APV)<br>6,3 (CGP)<br>7,5 (Bicuculline) | One way ANOVA |  | Overall | 2.277 x 10<sup>-5</sup>*** | F(3,27) = 12.72 |
|  |  |  |  | Bonferroni | Ctrl vs CNQX/APV | 1 |  |
|  |  |  |  |  | Ctrl vs CGP | 0.731 |  |
|  |  |  |  |  | Ctrl vs Bicuculline | 7.135 x 10<sup>-4</sup>*** |  |
| 2D | IPSCs – Mec/Atro | 8,6 | One way repeated measures ANOVA |  | Overall | 1.661 x 10<sup>-10</sup>*** | F(2,6) = 5454.97 |
|  |  |  |  | Tukey | Ctrl vs mec | 0.675 |  |
|  |  |  |  | Tukey | Ctrl vs atro | 2.472 x 10<sup>-9</sup>*** |  |
| 2D | IPSCs - TZP, AFDX, 4DAMP | TZP (5, 3),<br>AFDX(8, 3)<br>4DAMP(8, 3) | One way ANOVA |  |  | 5.807 x 10<sup>-10</sup>*** | F(3, 25) = 42.47 |
|  |  |  |  | Bonferroni | Ctrl vs TZP | 1 |  |
|  |  |  |  | Bonferroni | Ctrl vs AFDX | 1 |  |
|  |  |  |  | Bonferroni | Ctrl vs 4DAMP | 1.019 x 10<sup>-8</sup>*** |  |
| text | IPSCs: Male vs Female | Male (27, 10)<br>Female (24,8) | Two sample t-test |  | IPSC Frequency: Male vs. Female | 0.80 |  |
|  |  |  |  |  | IPSC Power: Male vs Female | 0.19 |  |
|  |  |  |  |  | IPSC Duration: Male vs Female | 0.72 |  |
| 4B | eNpHR: IPSC charge - baseline | CCK (9, 4),<br>PV (9, 3),<br>SOM (10, 4) | One-way ANOVA |  |  | 0.022* | F(2,25) = 4.48 |
|  |  |  |  | Bonferroni | CCK vs PV | 1 |  |
|  |  |  |  | Bonferroni | CCK vs SOM | 0.076 |  |
|  |  |  |  | Bonferroni | PV vs SOM | 0.035* |  |
| 4B | eNpHR: IPSC frequency - baseline | CCK (9, 4),<br>PV (9, 3),<br>SOM (10, 4) | One way ANOVA |  |  | 0.001*** | F(2,21) = 8.14 |
|  |  |  |  | Bonferroni | CCK vs PV | 1 |  |
|  |  |  |  | Bonferroni | CCK vs SOM | 0.003*** |  |
|  |  |  |  | Bonferroni | PV vs SOM | 0.011* |  |
| 4C | eNpHR: IPSC charge - ACh | CCK (9, 4),<br>PV (9, 3),<br>SOM (10, 4) | One-way ANOVA |  |  | 7.607 x 10<sup>-6</sup>*** | F(2,25) = 21.76 |
|  |  |  |  | Bonferroni | CCK vs PV | 0.001** |  |
|  |  |  |  | Bonferroni | CCK vs SOM | 6.13 x 10<sup>-6</sup>*** |  |
|  |  |  |  | Bonferroni | PV vs SOM | 0.027* |  |
| **4C** | eNpHR:<br>IPSC<br>frequency -<br>ACh | CCK (9, 4),<br>PV (9, 3),<br>SOM (10, 4) | One-way<br>ANOVA |  |  | 0.018* | F(2,21) = 4.92 |
|  |  |  |  | Bonferroni | CCK vs PV | 1 |  |
|  |  |  |  | Bonferroni | CCK vs SOM | 0.021* |  |
|  |  |  |  | Bonferroni | PV vs SOM | 0.056 |  |
| **4E** | eNpHR:<br>small and<br>large IPSCs | CCK (12, 4),<br>PV(16, 7),<br>SOM (13, 6) | Paired t-test | | CCK - large (average<br>IPSC frequency 2s prior<br>to light vs first 2s during<br>light | $5.123 \times 10^{-6***}$ | t(11) = 8.21 |
|  |  |  |  |  | PV - large | 0.274 | t(15) = 1.13 |
|  |  |  |  |  | SOM - large | 0.661 | t(12) = 0.66 |
|  |  |  |  |  | CCK - small | 0.001*** | t(11) = 4.16 |
| | | | | | PV - small | $4.88624 \times 10^{-4***}$ | t(15) = 4.43 |
|  |  |  |  |  | SOM - small | 0.672 | t(12) = 0.43 |
| **text** | PYR sIPSCs<br>across<br>different Cre<br>lines | CCK (9, 4),<br>PV (9, 3),<br>SOM (10, 4) | One way<br>ANOVA |  | Total IPSC frequency<br>baseline: CCK-, PV-,<br>SOM-Cre | 0.765 | F(2,25) = 0.269 |
|  |  | CCK (9, 4),<br>PV (9, 3),<br>SOM (10, 4) | One way<br>ANOVA |  | Total IPSC frequency<br>ACh: CCK-, PV-, SOM-Cre | 0.184 | F(2,22) = 1.829 |
|  |  | CCK (12, 4),<br>PV (16, 7),<br>SOM (13, 6) | One way<br>ANOVA |  | Large IPSC frequency<br>after ACh: CCK-, PV-,<br>SOM-Cre | 0.564 | F(2,38) = 0.582 |
|  |  | CCK (12, 4),<br>PV (16, 7),<br>SOM (13, 6) | One way<br>ANOVA |  | Small IPSC frequency<br>after ACh: CCK-, PV-,<br>SOM-Cre | 0.256 | F(2,38) = 1.414 |
| **6B** | CCK, PV,<br>and SOM IN<br>properties:<br>resting<br>membrane<br>potential | CCK (18, 7),<br>PV (132, 45),<br>SOM (48, 17) | One-way<br>ANOVA | | | $6.159 \times 10^{-13***}$ | F(2,195) = 32.59 |
|  |  |  |  | Bonferroni | PV vs CCK | 0.004** |  |
|  |  |  |  | Bonferroni | SOM vs CCK | 0.208 |  |
| | | | | Bonferroni | SOM vs PV | $8.088 \times 10^{-13***}$ | |
| **6B** | CCK, PV,<br>and SOM IN<br>properties:<br>membrane<br>resistance | CCK (18, 7),<br>PV (132, 45),<br>SOM (48, 17) | One-way<br>ANOVA | | | $1.699 \times 10^{-26***}$ | F(2,195) = 81.68 |
| | | | | Bonferroni | PV vs CCK | $1.038 \times 10^{-7***}$ | |
|  |  |  |  | Bonferroni | SOM vs CCK | 0.076 |  |
| | | | | Bonferroni | SOM vs PV | $1.628 \times 10^{-25***}$ | |
| **6F** | CCK firing<br>with 4DAMP | 4, 3 | Paired t-test | | Ctrl vs 4DAMP | $9.701 \times 10^{-4***}$ | t (3) = 13.05 |
| **6G** | CCK depol.<br>with 4DAMP | 4, 3 | Paired t-test |  | Ctrl vs 4DAMP | 0.016* | t(3) = 4.97 |
| **7B** | Theta MPO<br>before and<br>after light | 38, 13 | Paired t-test | | Before light vs After light | $2.283 \times 10^{-5***}$ | t(37) = -4.84 |
| **7C** | Theta MPO<br>with bic | 8,5 | One-way<br>repeated<br>measures | | | $5.243 \times 10^{-4***}$ | F(2,6) = 34.20 |
| | | | | Tukey | After light vs Before light | $5.83 \times 10^{-4***}$ | |
| | | | | Tukey | After + bic vs Before light | $2.58 \times 10^{-4***}$ | |
| **7D** | Theta MPO with WIN | 9,3 | One-way repeated measures | | multivariate | 0.032* | $F(2,7) = 5.88$ |
|  |  |  |  | Tukey | Before light vs After light | 0.001** |  |
|  |  |  |  | Tukey | Before light vs After light + WIN | 0.95675 |  |
|  |  |  |  | Tukey | After light vs After light + WIN | 0.003** |  |
| **7F** | Cell attached firing | Ctrl (11,7), Atro (4,3) | Two sample t-test | | Firing frequency after light: Ctrl vs Atro | 0.022 | $t(14) = 2.56$ |
| **8B** | MPO synchrony | 23,12 | Paired t-test | | Before light vs After light | $6.25 \times 10^{-5***}$ | $t(22) = -4.92$ |
| **8F** | Firing synchrony | Control (10,8), Picro (3,3) | Two sample t-test | | Control vs Picro | 0.015* | $t(11) = 1.58$ |

**Supplemental Figure 1.**
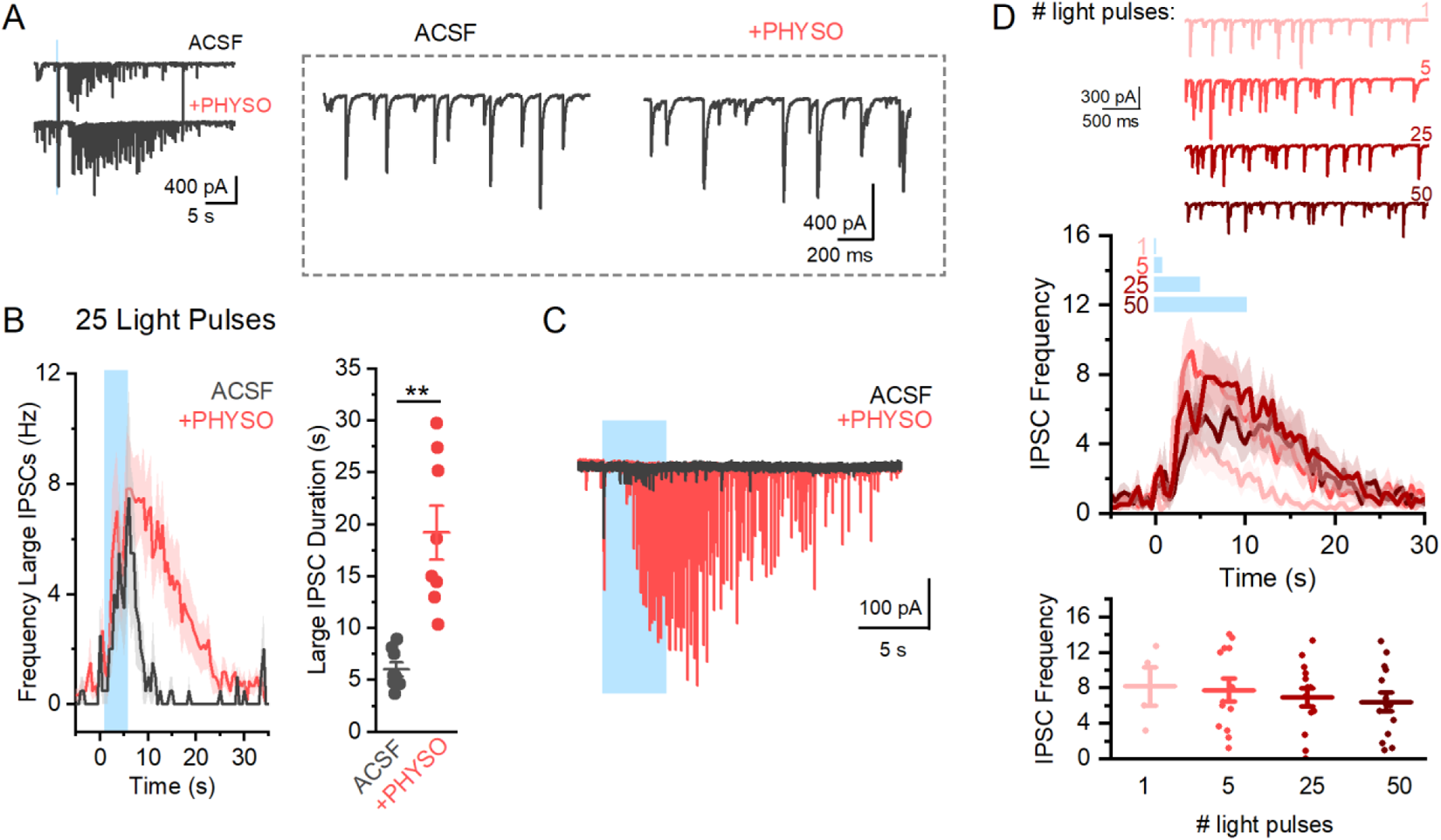
Increasing cholinergic stimulation and the effect of physostigmine on BLA IPSCs. **A.** Sample trace from a BLA PYR showing large IPSCs in response to cholinergic stimulation in ACSF and PHYSO (1 μM) that shows similar IPSC frequency across conditions (inset, right). **B.** Comparing the time course and frequency of large IPSCs induced by ACh (5 Hz, 5 s) in ACSF and PHYSO conditions shows that the frequency of IPSCs induced by ACh does not change with the addition of PHYSO (IPSC frequency following light in cells where large IPSCs were induced: ACSF = 7.0 ± 1.71 Hz, n = 4, N = 3; +PHYSO = 7.37 ± 1.18 Hz; n = 12, N = 7, two sample t-test, p = 0.87, t(14) = −0.16). Instead, the duration of the IPSCs was significantly increased by ACh (additional set of PYRs comparing only cells where large IPSCs induced in ACSF and PHYSO conditions, large IPSC duration: ACSF = 6.05 ± 0.66 s; PHYSO = 19.20 ± 2.57 s, n = 8, N = 5, paired t-test, p = 0.003, t(7) = −0.16). **C.** Representative trace showing a BLA PYR where IPSCs were not induced in ACSF conditions but could be produced in PHYSO. **D.** Representative traces of large IPSCs induced by ACh after 1, 5, 25, or 50 pulses of light show that the frequency of IPSCs does not change across stimulation parameters (bottom graph, average IPSC frequency following 1, 5, 25, and 50 pulses of light are not significantly different, one way ANOVA, p = 0.80, F(3) = 0.33).

**Supplemental Figure 2.**
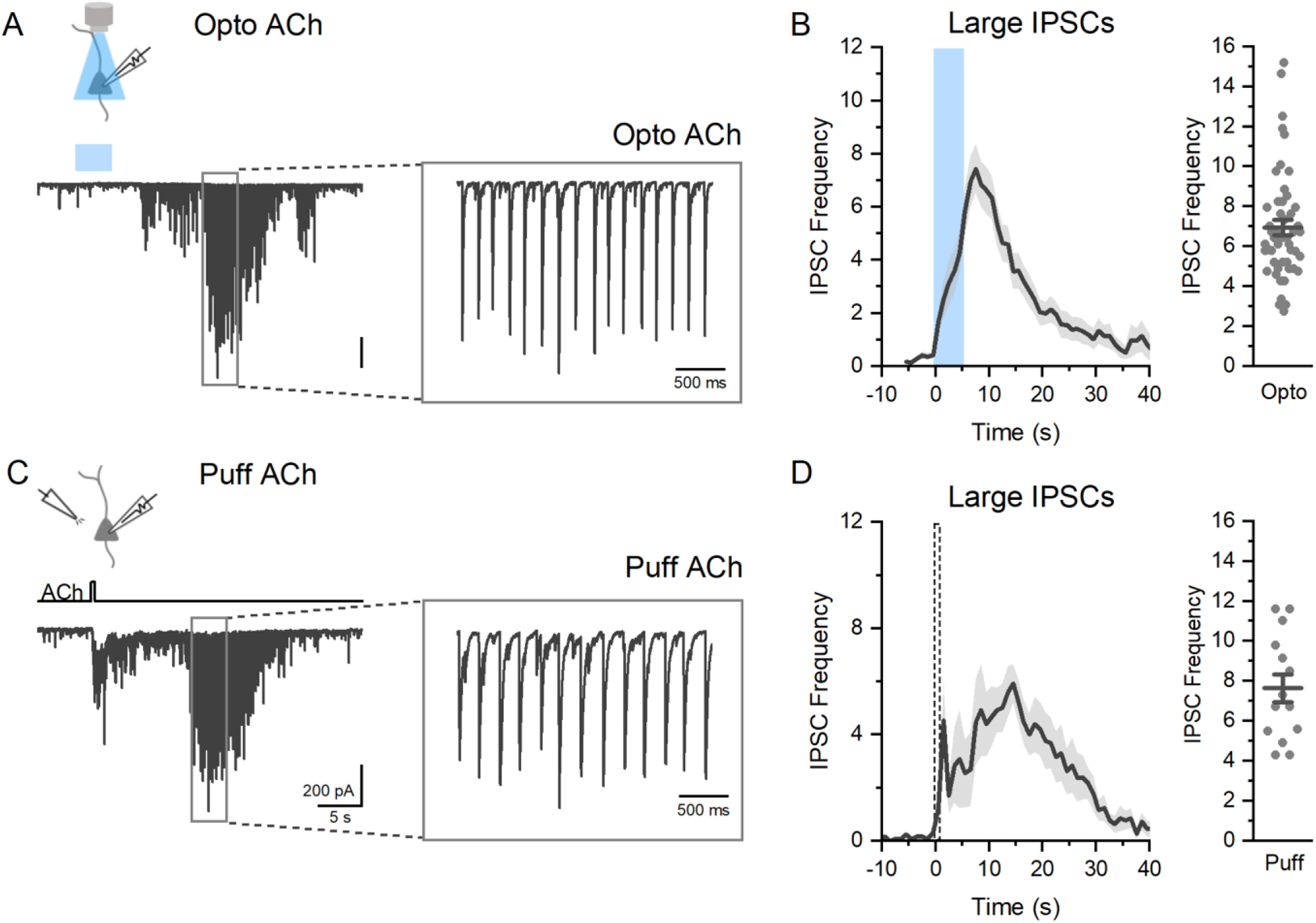
Comparing the effects of optogenetically released acetylcholine to focal puff application of acetylcholine. **A.** Sample waveforms from two separate BLA PYRs showing large rhythmic IPSCs (inset) in response to optogenetically released ACh (top) or focal puff application of ACh (bottom: 500 ms duration, 1 mM ACh, within 100 microns of recorded PYR). **B.** Comparing the frequency of large IPSCs over time shows that both optogenetically released ACh and focal puff application of ACh induce prolonged increases in IPSCs that peak in the theta frequency seconds after the cholinergic stimulus. Comparisons of large IPSC frequency shows no difference between the different stimulations (optogenetically released ACh = 7.63 ± 0.71 Hz, n = 51, N = 18; focal puff ACh = 6.94 ± 0.38 Hz, n = 14, N = 12, two sample t-test, p = 0.39, t(63) = 0.85) indicating they are inducing similar networks effects in BLA brain slices.

**Supplemental Figure 3.**
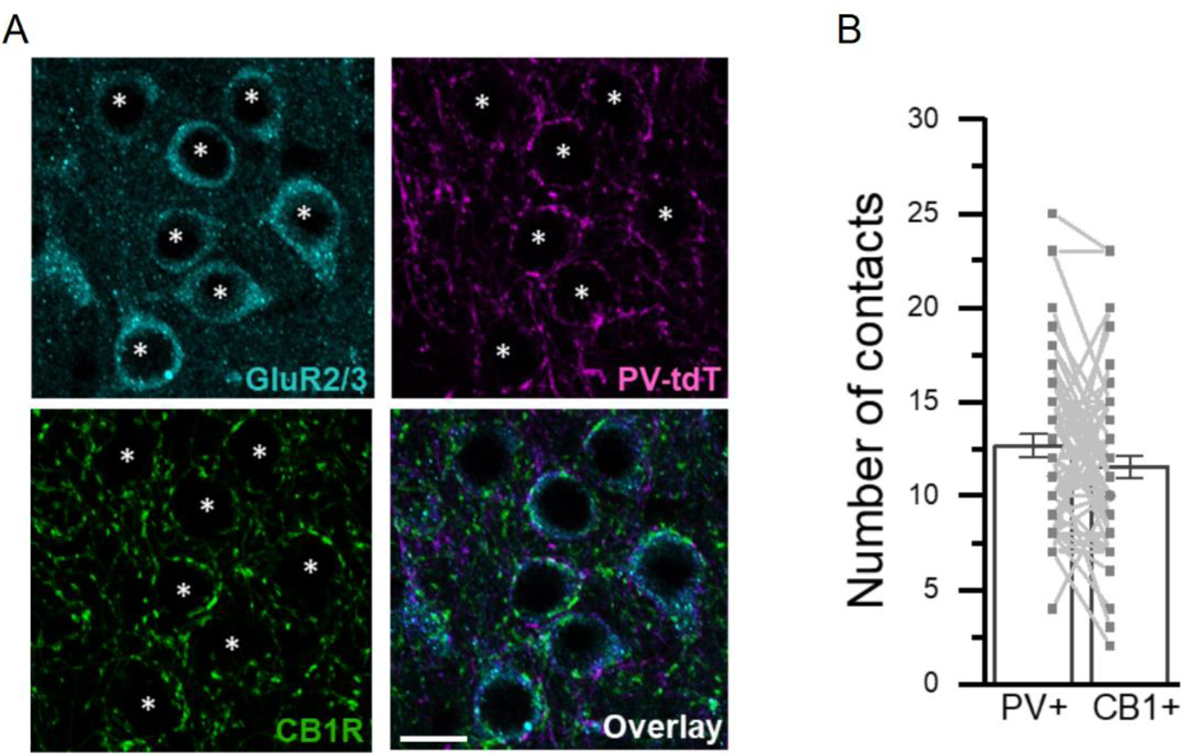
Equal contacts on BLA PYRs by PV and CB1+ synapses. **A**. Representative image of BLA PYRs (expressing GluR2/3, cyan) surrounded by PV (magenta) and CB1+ (green) contacts. Scale bar 25 μm. **B.** Comparing the total number of contacts on BLA PYR cell bodies shows no difference between PV and CB1+ contacts (# of contacts: PV = 12.64 ± 0.62; CB1 = 11.53 ± 0.59; n = 46 BLA PYRs, N = 3, paired t-test, p = 0.09, t(55) = 1.71).

**Supplemental Figure 4.**
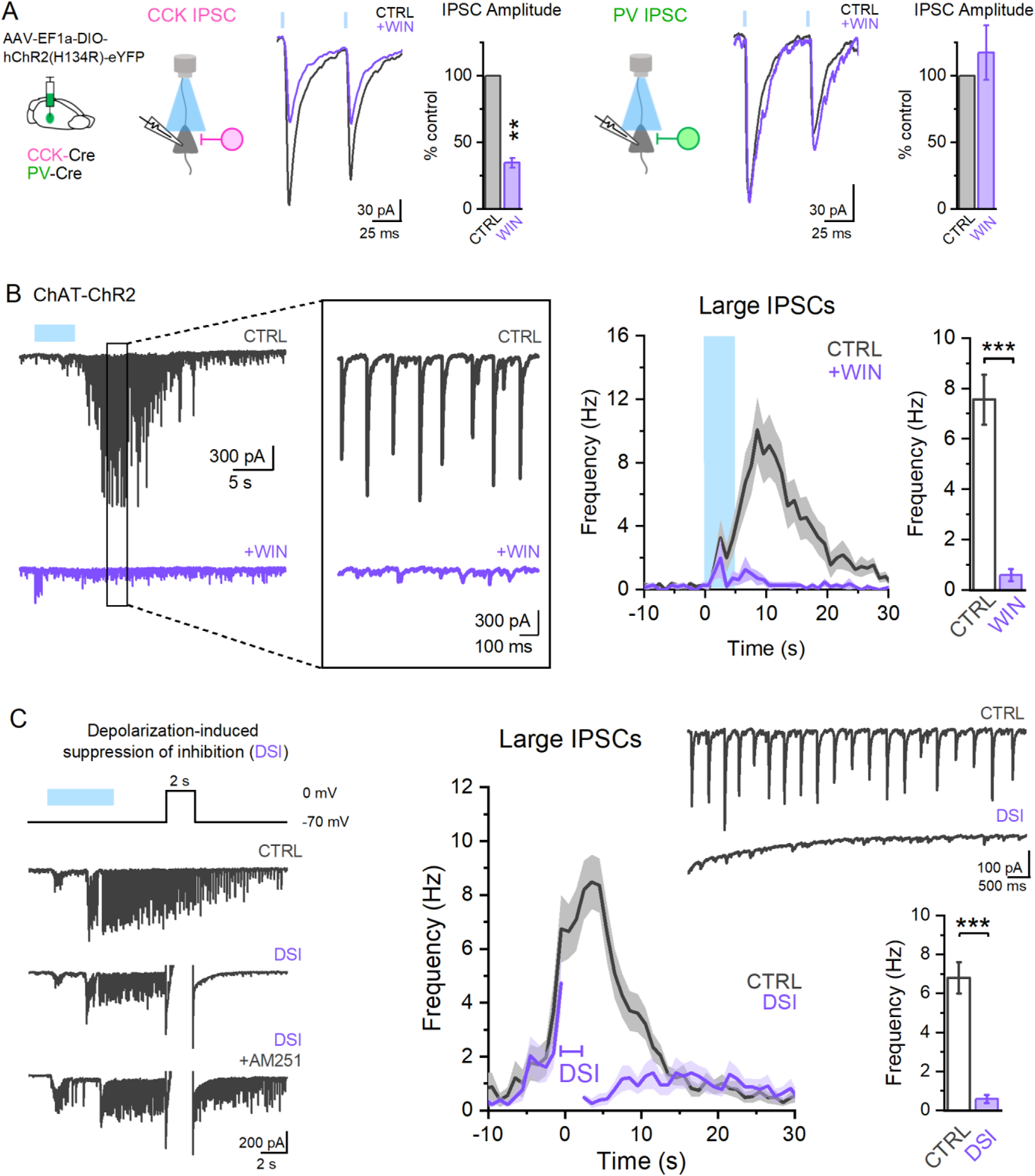
Cannabinoid receptor 1 (CB1R) expressing CCK interneurons underly ACh-induced large IPSCs in BLA PYRs. **A.** In CCK-Cre and PV-Cre mice AAV5-EF1a-DIO-hChR2(H134R)-eYFP was injected into the BLA to allow for selective activation of CCK and PV INs in brain slice preparations. Application of WIN 55,212-2 (2 μM) reduced the amplitude of CCK-evoked IPSCs (WIN = 0.35 ± 0.04 of control IPSC amplitude; n = 4, N = 3, paired t-test, p = 0.003, t(2) = 17.61) but not PV IPSCs (WIN = 1.17 ± 0.21 of control IPSC amplitude; n = 3, N = 3; paired t-test, p = 0.48, t(2) = −0.85). **B.** Sample recording from BLA PYR showing large IPSCs in response to ACh stimulation in control conditions that are abolished by application of WIN in the bath. Measurements of large IPSC frequency show that these events are blocked by WIN (large IPSC frequency after ACh: CTRL = 7.56 ± 0.99 Hz, +WIN = 0.59 ± 0.25 Hz; n = 11, N = 5, p = 6.05 x 10^−5^, t(10) = 6.60). **C.** Additionally, depolarization-induced suppression of inhibition (DSI) in a PYR recording (holding at 0mV for 2s) could also block large IPSCs after ACh stimulation that could be reversed by AM251 (1 μM, n = 5, N = 4), indicating an involvement of CB1Rs. Measuring large IPSC frequency in control conditions and after DSI shows this significant reduction (large IPSC frequency after ACh: CTRL = 6.82 ± 0.81 Hz; after DSI = 0.58 ± 0.20 Hz; 23 cells, 11 animals; paired t-test, p = 1.11 x 10^−7^, t(22) = 7.70).

**Supplemental Figure 5.**
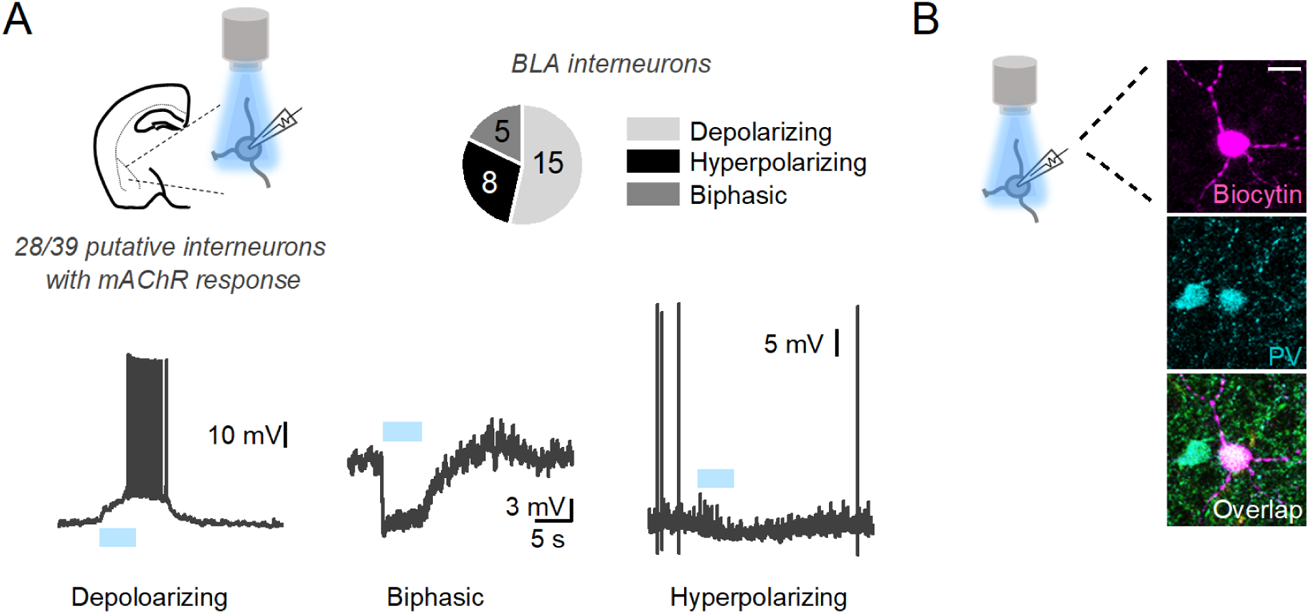
Endogenous acetylcholine release on putative BLA interneurons. **A.** In ChAT-ChR2 brain slice preparations, recordings from putative INs were made as determined by electrophysiological parameters. Of 39 putative INs across 16 animals, 28 cells exhibited slow muscarinic responses in response to ACh stimulation. Putative IN muscarinic responses could be classified as either depolarizing, biphasic, or hyperpolarizing and looked similar in nature to the response profiles observed to focal puff ACh (Figure 6B). **B.** In 4 experiments, post-hoc labeling of the biocytin-filled recorded cell co-labelled with PV antibody, indicating recordings from PV INs, that showed depolarizing responses, consistent with the response of these cells to focal ACh application. Scale bar 25 μm.

**Supplemental Figure 6.**
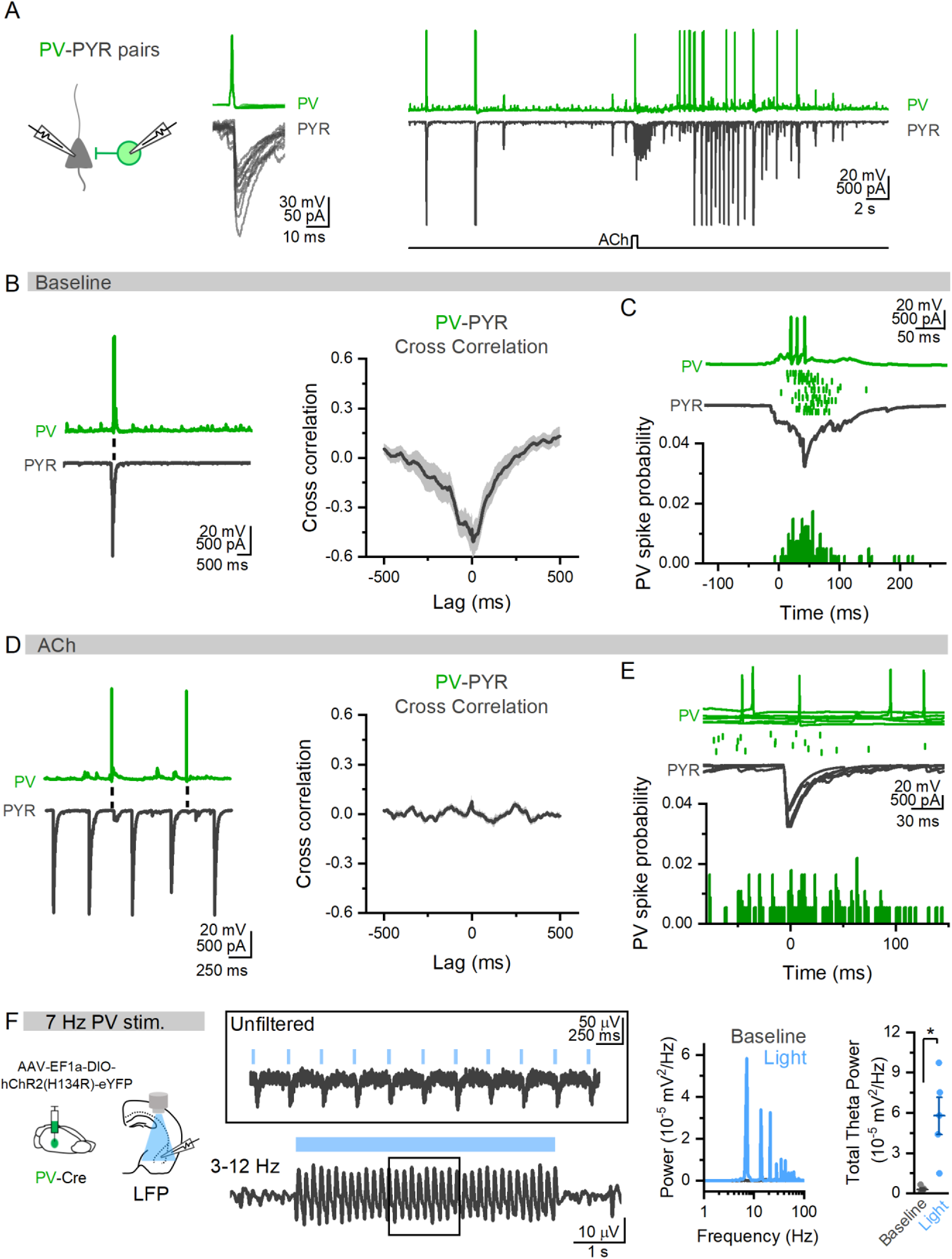
BLA PV interneurons align with large network activity in baseline but not during ACh states. **A.** Example paired recording of a synaptically connected PV IN and PYR in the BLA (left) in baseline and after puff application of ACh (right). **B.** In same PV-PYR pair, sample trace of baseline network activity in the absence of ACh showing PV firing and large IPSCs in BLA PYR. Cross correlation of these baseline network events shows high degree of cross correlation (n = 5 pairs, N = 4) **C.** Zoomed in traces showing PV firing in relation to PYR network event (green dots represent action potentials across multiple trials from the same PV IN), shows these events are tightly aligned to PYR activity in baseline conditions (bottom, total of 100 PV spikes from 5 PV-PYR pairs, 4 animals) **D.** In the same PV-PYR pair, traces show action potential firing in PV INs does not align with PYR IPSCs in the presence of ACh. Cross correlation of PV and PYR membrane potentials after ACh show low cross correlation (13 pairs, N = 6). **E.** Time-locked overlay of 5 individual large IPSCs after ACh in the PYR (bottom) and action potentials in PV IN (top, dots underneath trace represent PV action potentials across 11 IPSCs) highlights that these events do not align (bottom, probability of PV action potential in relation to the start of large IPSC in PYR: total of 107 PV spikes from 13 PV-PYR pairs, 6 animals). **F.** When ChR2 is expressed in PV INs in BLA slices, light stimulation at 7 Hz results in robust oscillations of the LFP (traces from example slice and corresponding power spectrum, right). Comparing total theta power (3-12 Hz) of LFP in baseline and during light stimulation of PV INs shows induction of theta oscillations by PV INs (total theta power (10^−5^ mV^2^/Hz): baseline = .28 ± .10; light = 5.78 ± 1.40; n = 5, N = 3, paired t-test, p = 0.016, t(4) = −4.01).

**Supplemental Figure 7.**
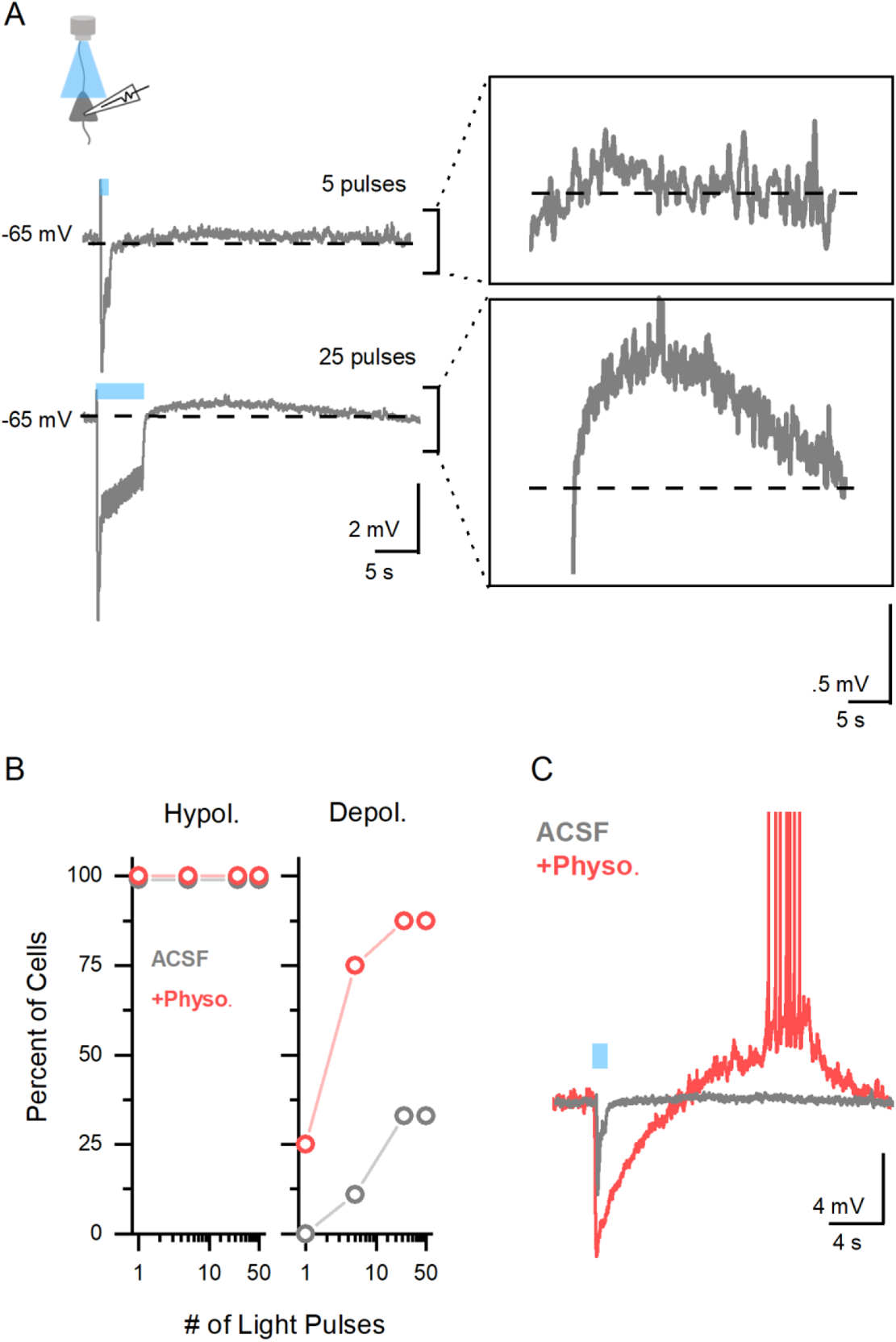
Large, but not small, cholinergic stimuli are required to depolarize BLA PYRs. **A.** Sample trace from recording of BLA PYR showing an initial hyperpolarization during light stimulation to both 5 and 25 pulses of light stimulation, but only the 25 pulse stimulation evoked a depolarizing response (zoomed in, inset right). **B**. Plotting the proportion of BLA PYRs that show a hyperpolarizing response to increasing amounts of ACh stimulation show that initial hyperpolarizing responses are evoked in all cells (left) while prolonged depolarizing responses require increasing stimulation or the presence of physostigmine (PHYSO, 1 uM, right) (n = 9, N = 3). **C.** A representative recording of a BLA PYR where addition of physostigmine caused a small acetylcholine stimulus that does not induce PYR depolarization to evoke a large PYR depolarization and action potential firing.

**Supplemental Figure 8.**
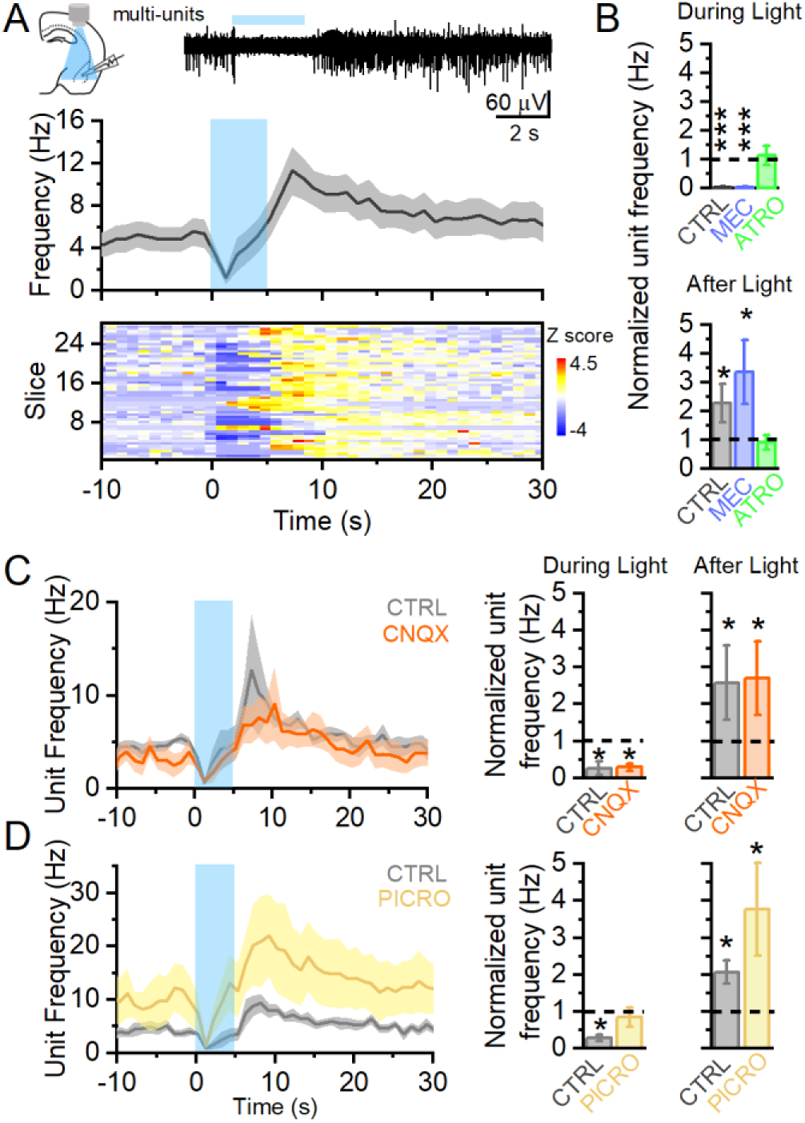
Acetylcholine induces biphasic changes in BLA network activity via muscarinic receptors. **A.** Multi-unit recordings show an initial suppression of unit frequency by cholinergic stimulation (5 Hz, 5s) that is then followed by a robust increase in unit activity (normalized unit frequency: during light = 0.03 ± 0.03 of baseline; paired t-test, p = 0.001; after light = 2.26 ± 0.66 of baseline; paired t-test = 0.047; n = 27, N = 15). Raster plot of unit activity and heat map of unit frequency show these bidirectional changes (bottom). **B.** Application of mecamylamine (Mec,10 μM) did not block the changes in unit activity by ACh during light or after light, but application of atropine (Atro, 5 μM) blocked any changes by acetylcholine (5 μM) (Unit frequency during light: CTRL = 0.03 ± 0.03 of baseline, one way ANOVA repeated measures, post hoc Tukey p = 1.65 x 10^−4^; Mec. = 0.04 ± 0.03 of baseline; p = 1.88 x 10^−4^; Atro. = 1.14 ± 0.33 of baseline, p = 0.99; n = 6, N = 5). **C.** Unit activity in response to cholinergic stimulation was not affected by CNQX (20 μM) (unit frequency during light: control = 0.26 ± .17 of baseline; CNQX = 0.29 ± 0.09; paired t-test, p = 0.83; n = 5, N = 4). **D.** Application of picrotoxin to block GABA_A_ inhibition enhanced the frequency of unit activity during and after light stimulation (unit frequency during light: control = 0.28 ± 0.09 of baseline, Picro = 0.85 ± 0.26 of baseline; after light: control = 2.06 ± 0.31, Picro = 3.77 ± 1.26, n = 6, N = 4).

## Notes

### Competing Interest Statement

The authors have declared no competing interest.

